# Bsp22 protein polymerization drives dynamic adaptation of the *Bordetella* type III secretion system injectisome

**DOI:** 10.1101/2024.03.04.583273

**Authors:** Ivana Malcova, Jana Kamanova

**Affiliations:** Laboratory of Infection Biology, Institute of Microbiology of the Czech Academy of Sciences, Videnska 1083, 142 20 Prague 4, Czech Republic

## Abstract

Many gram-negative bacteria are equipped with a type III secretion system (T3SS) injectisome, enabling the direct translocation of effector proteins from the bacterial cytoplasm into host cells. In the case of *Bordetella bronchiseptica*, a respiratory pathogen of diverse mammals, the T3SS injectisome exhibits a unique needle tip filament formed by the Bsp22 protein, essential for bacterial persistence in mice. Here, we used *B. bronchiseptica* and *B. pertussis* with in-frame insertions of short peptide tags into the Bsp22 to elucidate its formation and characteristics via super-resolution imaging with fluorophore-labeled nanobodies and biarsenic probes. We report that on abiotic surfaces, Bsp22 forms flexible filaments that protrude up to several µm from bacterial cells. In these conditions, Bsp22 filaments grow continuously without any apparent growth control, with Bsp22 subunits being added at the distal end. Remarkably, during host cell infection, the growth of Bsp22 filaments is constrained, accompanied by downregulated Bsp22 protein production. When infecting HeLa cells, some Bsp22 filaments form a short physical bridge between the bacterium and the host cell surface. In nasal epithelium, where *B. bronchiseptica* colonization is specifically restricted to ciliated cells, Bsp22 filaments align parallel to the cilia, pointing towards the cell apical surface. These results highlight the context-specific and environment-influenced dynamic modulation of the length and orientation of *Bordetella* needle tip filament, uncovering the adaptability of the *Bordetella* T3SS injectisome in response to conditions encountered during host cell infection.

## Introduction

The classical *Bordetella* species, specifically *B. bronchiseptica, B. pertussis,* and *B. parapertussis* are respiratory pathogens that colonize the ciliated epithelial cells of the mammalian respiratory tract. *B. bronchiseptica*, infects a variety of mammals, leading to diverse pathologies. These include persistent, often asymptomatic respiratory infections, as well as acute diseases such as kennel cough in dogs, and bronchopneumonia and atrophic rhinitis in piglets [1]. The strictly human-adapted *B. pertussis* and the human-adapted lineage of *B. parapertussis*_HU_, on the other hand, are responsible for causing whooping cough, also known as pertussis, which remains a significant concern for public health and one of the least controlled vaccine-preventable infectious diseases [2–4]. It is believed that *B. pertussis* and *B. parapertussis*_HU_ evolved independently from a *B. bronchiseptica*–like ancestor, sharing highly identical genetic content, including genes that encode the type III secretion system (T3SS) injectisomes [5–7].

The T3SS injectisomes function as syringe-like nanomachines embedded in the bacterial envelope, facilitating the direct translocation of effector proteins from the bacterial cytosol into the host cell cytoplasm. The translocation process involves recruiting secreted effector proteins to the cytoplasmic sorting platform, loading them into the inner membrane export apparatus, and subsequently transporting them through a continuous channel within the injectisome basal body and the extracellular needle filament. The needle filament is capped by a tip complex, connecting it to the translocon pore in the host cell membrane, thus creating a complete channel [8, 9]. These nanomachines likely arose by exaptation of the bacterial flagellum, which was followed by their rapid diversification and adaptation to different host cells and bacterial lifestyles [10, 11].

The prototypical injectisome, as encoded in pathogens such as *Salmonella*, *Shigella*, and *Yersinia,* is characterized by a needle filament measuring 45-80 nm in length, capped by a ring-shaped pentameric tip complex composed of the tip protein [12–16]. Interestingly, in enteropathogenic *Escherichia coli* (EPEC), this pentameric tip complex is replaced by a tip filament, an extended structure connected to the needle, formed through the polymerization of a tip protein EspA [17–20]. Similarly, in *B. bronchiseptica*, immunoelectron microscopy has shown that a tip protein Bsp22 forms surface appendages [21]. Due to the absence of significant sequence similarity to EspA and the pentameric T3SS-tip complex protein families SipD/IpaD (*Salmonella* and *Shigella* spp.) and LcrV/PcrV (*Yersinia* and *Pseudomonas* spp.), it has been proposed that Bsp22 defines a novel T3SS-tip complex family [21, 22].

Bsp22 is the most abundantly secreted T3SS substrate of *B. bronchiseptica*. Its role in the *Bordetella* injectisome tip complex is evidenced by the fact that Bsp22 inactivation does not block T3SS-mediated secretion into the medium but does block the translocation of the cytotoxic BteA effector into host cells, thereby hindering T3SS-mediated cytotoxicity [21, 23, 24]. Additionally, similar to *Bordetella* T3SS ATPase BscN, Bsp22 is crucial for the persistence of *B. bronchiseptica* in mouse tracheas [23]. Moreover, while immunization of mice with Bsp22 does not confer protection against an intranasal challenge with *B. pertussis*, it does lead to a significant reduction in colonization by *B. bronchiseptica* [21, 25].

Although the significance of Bsp22 filaments is recognized, there remains a gap in our understanding regarding their formation and dynamics during *Bordetella* infection, as well as their potential similarities or differences compared to EspA filaments of EPEC. Therefore, the aim of this study was to visualize Bsp22 filaments on abiotic surfaces and during host cell infection using super-resolution fluorescence imaging. This exploration is crucial for comprehending how classical *Bordetella* species interact with their hosts, and it also broadens our understanding of the assembly and function of different families of T3SS injectisomes.

## Results

### The Bsp22 protein polymerizes into flexible filaments that intertwine on the abiotic surface of a glass coverslip

To enable super-resolution fluorescence imaging, we first investigated whether Bsp22 would tolerate the insertion of a short peptide SPOT-tag recognized by a commercially available nanobody without affecting Bsp22 function in the *Bordetella* injectisome. To select a suitable insertion site, we examined the three-dimensional structure of Bsp22 from *B. bronchiseptica* RB50 predicted by AlphaFold. The structure revealed a long N-terminal α-helix (I26-A74) connected to a C-terminal α-helix (Y144-M204) by a mixed α/β-region, with a surface-exposed unstructured loop (T135-T143) (Figure 1A). Given that a similar surface-exposed loop in the EspA protein was previously demonstrated to tolerate insertions [26, 27], we opted to insert the SPOT-tag flanked by GSSG linkers between amino acid positions M138 and A139 of Bsp22, resulting in a *B. bronchiseptica* RB50 derivative, *Bb bsp22*^SPOT^. To assess the injectisome functionality of the *Bb bsp22*^SPOT^, we examined its ability to induce T3SS-dependent cell cytotoxicity mediated by injection of the type III effector protein BteA. As shown in Figure 1B, *Bb bsp22*^SPOT^ exhibited cell cytotoxicity comparable to that of the wild-type *Bb* strain. After a 3-hour infection, both strains induced lysis of 50% and 80% of HeLa cells at a multiplicity of infection (MOI) of 5:1 and 50:1, respectively. In contrast, infection with RB50 derivatives harboring in-frame deletions of the open reading frames for Bsp22 (*Bb* Δ*bsp22*) or the type III effector BteA (*Bb* Δ*bteA*) did not induce cell lysis. These data demonstrate that the Bsp22 protein can tolerate the insertion of a short peptide tag between amino acids M138 and A139.

**Figure 1.**
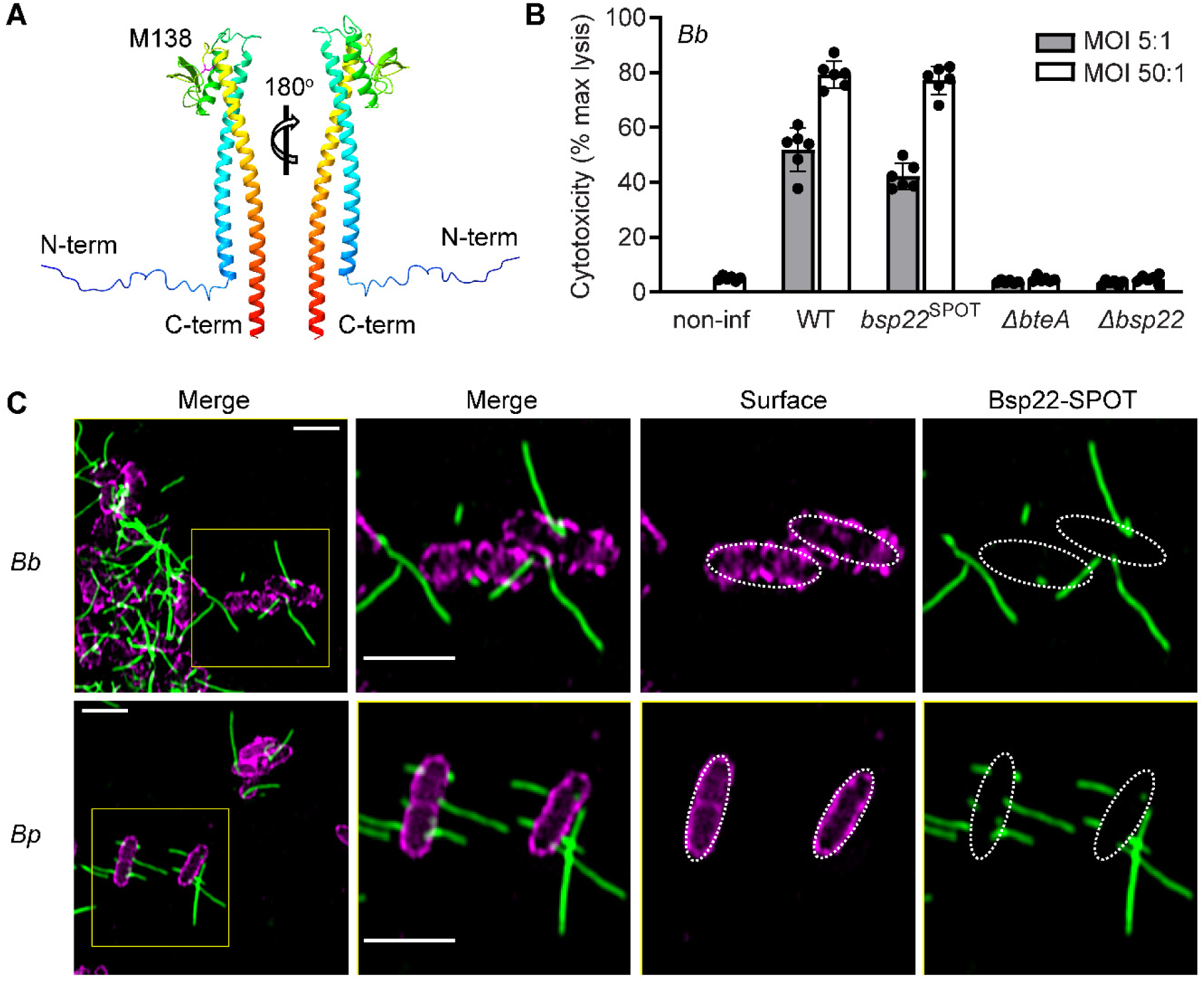
The Bsp22 protein polymerizes into flexible filaments that intertwine on the abiotic surface of a glass coverslip. (A) Ribbon representation of the monomeric AlphaFold-predicted structure of Bsp22 from B. bronchiseptica RB50. The unstructured loop of the mixed α/β-region, with M138 highlighted, is shown. This figure was generated by UCSF ChimeraX. (B) The functionality of the *Bordetella* injectisome is not affected by the insertion of the SPOT-tag between amino acids M138 and A139 of Bsp22. T3SS-dependent cytotoxicity of the indicated *B. bronchiseptica* strains towards HeLa cells was measured at a multiplicity of infection (MOI) of 5:1 and 50:1 as the amount of lactate dehydrogenase (LDH) released 3 h post-infection. Mean values ± SD from two biological replicates with three technical replicates each are shown. (C) Bsp22 forms filaments on the surface of *B. bronchiseptica* and *B. pertussis*. Cells of *B. bronchiseptica* RB50 (*Bb*) or *B. pertussis* B1917 (*Bp*) carrying *bsp22*^SPOT^ were cultured on coverslips for 3 h. After fixation, Bsp22 (green) was visualized with Spot-label ATTO488, while the bacterial outer surface (magenta) was labeled with a rabbit anti-*Bordetella* serum followed by anti-rabbit IgG-DyLight 405 conjugate. The SIM images show a single focal plane and are representative of three independent experiments. Scale bars, 2 μm.

Next, we employed structured illumination microscopy (SIM) to examine the Bsp22 appendages in the *Bb bsp22^SPOT^* strain. Cells of *Bb bsp22^SPOT^*were spotted onto the abiotic surface of glass coverslips and cultured for 3 hours, followed by fixation and labeling of Bsp22 using an anti-SPOT nanobody conjugated to ATTO488, Spot-label ATTO488. As shown in Figure 1C and Figure S1, Bsp22 polymerized into filaments that extended in different directions from *Bb* cells and had varying lengths. These filaments were found in very low numbers per bacterial cell, ranging from none to a few, with some reaching lengths of over two μm. We also observed that these filaments intertwine and twist, indicating their flexibility. Importantly, similar Bsp22 filaments of different lengths, twisting in different directions, were also produced by *B. pertussis* B1917 derivative, *Bp bsp22*^SPOT^, in which we introduced the SPOT-tag at the same position within Bsp22, that is, in between amino acids M138 and A139 (Figure 1C). Thus, both classical *Bordetella* species, *B. bronchiseptica*, which causes chronic respiratory infections in a variety of mammals, and *B. pertussis*, an exclusively human pathogen causing whooping cough, produce long and flexible Bsp22 filaments that intertwine on abiotic surfaces, such as glass coverslips, during bacterial cell growth. Additionally, Bsp22 tolerates the insertion of short peptide tags, allowing for the peptide display on the surface of Bsp22 filaments, a feature similar to the EspA protein.

### Growth of Bsp22 filaments is continuous, with Bsp22 monomers being added at the distal filament end

When visualizing Bsp22, we observed filaments of different lengths, suggesting a lack of growth control during their assembly. To analyze the dynamics of filament formation, we cultured *Bb bsp22*^SPOT^ on glass coverslips and fixed them at different time points: 0.5, 1, 2, 3, and 6 hours of incubation. When the bacteria were fixed only 0.5 hour after spotting, they exhibited short filaments. We also noticed some filaments that were torn from the bacterial surface, which was likely due to shaking of the initial bacterial culture or shear force caused by pipetting (Figure 2A and Figure S2). As the incubation continued, the length of Bsp22 filaments increased, demonstrating their continual growth on abiotic surface. Interestingly, 6 hours after spotting, filaments even formed a cohesive network, as shown in Figure S2.

**Figure 2.**
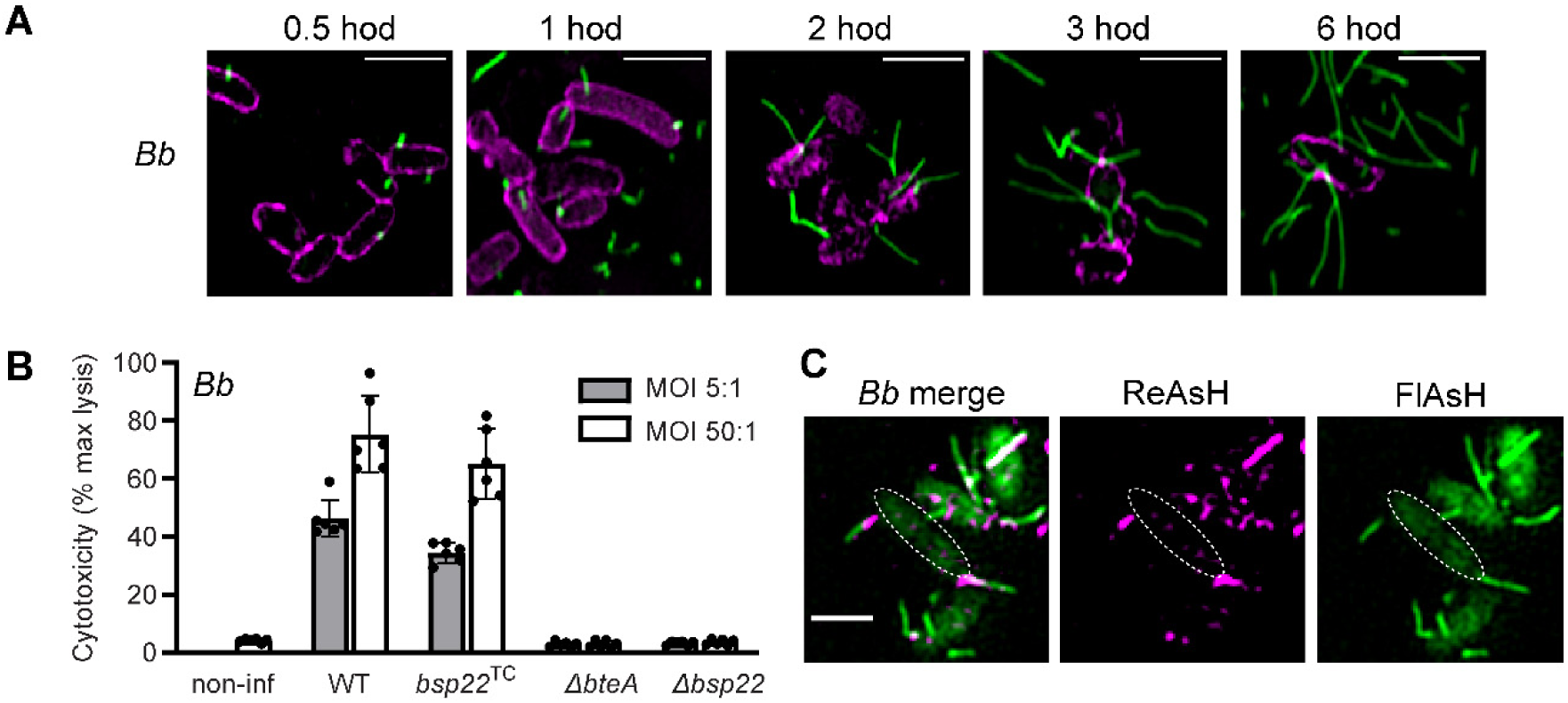
Growth of Bsp22 filaments is continuous, with Bsp22 monomers being added at the distal filament end. (A) Growth of Bsp22 filaments is continuous. Cells of *Bb bsp22*^SPOT^ incubated on coverslips were fixed at the indicated time points. Staining was performed as described in Figure 1C legend. Bsp22, green; *Bb* outer surface, magenta. The SIM images of a single focal plane are representative of 3 independent experiments. Scale bars, 2 μm. (B) Tagging of Bsp22 by TC-tag does not affect the functionality of the Bordetella injectisome. T3SS-dependent cytotoxicity of the indicated Bb strains towards HeLa cells was measured as described in Figure 1B legend. Mean values ± SD from two biological replicates with three technical replicates each are shown. (C) Bsp22 monomers are added at the distal filament end. Cells of *Bb bsp22*^TC^ were sequentially labeled with ReAsH (magenta) followed by FlAsH (green) in a pulse-chase experiment and analyzed by fluorescence microscopy. The SIM images of fixed cells show a single focal plane and are representative of 3 independent experiments. Scale bars, 2 μm.

We next investigated the incorporation site of newly added Bsp22 subunits within the filament by employing an optimized tetracysteine tag (TC-tag) designed for binding biarsenical dyes [28]. The SPOT-tag was substituted for the TC-tag positioned between amino acids M138 and A139 of Bsp22 in *Bb* RB50, generating the *Bb bsp22*^TC^ strain. Importantly, this strain maintained T3SS-dependent cytotoxicity (Figure 2B), confirming the continued functionality of the T3SS injectisome. To assess the addition of Bsp22 subunits to filaments, we performed a pulse-chase experiment using two different biarsenical dyes, fluorescein arsenical hairpin (FlAsH) and resorufin arsenical hairpin (ReAsH), with *Bb bsp22*^TC^ cells in solution. The bacteria were labeled with ReAsH, washed, and then incubated with FlAsH. As illustrated by a representative image of double-labeled cells in Figure 2C, a one-hour incubation with FIAsH was sufficient to clearly detect the growing Bsp22 filament. Moreover, the ReAsH dye, applied during the pulse, stained the proximal segment of the filament, while the FIASH dye added during the subsequent chase labeled the distal end of the growing filament. These data demonstrate that formation of Bsp22 filaments on the abiotic surface or in a solution is an ongoing process. Moreover, newly incorporated Bsp22 subunits are added at the distal end of the growing filament, which is reminiscent of the formation of flagellar filaments.

### Bsp22 filaments are less abundant than injectisome rings and their presence is not influenced by calcium regulation

In further exploration of Bsp22 filament biogenesis, we examined the influence of physiological concentrations of Ca^2+^ ions, known to regulate T3SS effector protein secretion in *B. bronchiseptica* [24]. As depicted in Figure 3 and Figure S3A, the addition of 2 mM Ca^2+^ ions during the cultivation of *B. bronchiseptica* had no effect on the distribution pattern of Bsp22 filaments on cells. This observation is consistent with previous findings that Ca^2+^ ions have no effect on the total amount of Bsp22 protein in the culture supernatant [24]. Interestingly, as also shown in Figure 3, when we visualized the Bsp22 protein together with the inner-membrane ring component BscD of the injectisome by tagging its cytoplasmic N-terminus with the fluorescent protein mNeonGreen (mNG), it became evident that Bsp22 filaments are less abundant than intracellular BscD foci. Importantly, the functionality of the injectisome was not affected by the tagging of the BscD protein, as previously reported [24]. This was confirmed by the fact that the resulting strain, *Bb bsp22*^SPOT^ / *mNG-bscD*, induced the same T3SS-dependent cytotoxicity as the wild-type *Bb* strain or *Bb bsp22*^SPOT^ derivative (see Figure S3B). In summary, the growth of Bsp22 filaments was not regulated by Ca^2+^ ions. Regardless of the Ca^2+^ concentration, Bsp22 filaments remained in lower numbers compared to membrane injectisome components, with only a few per bacterial cell.

**Figure 3.**
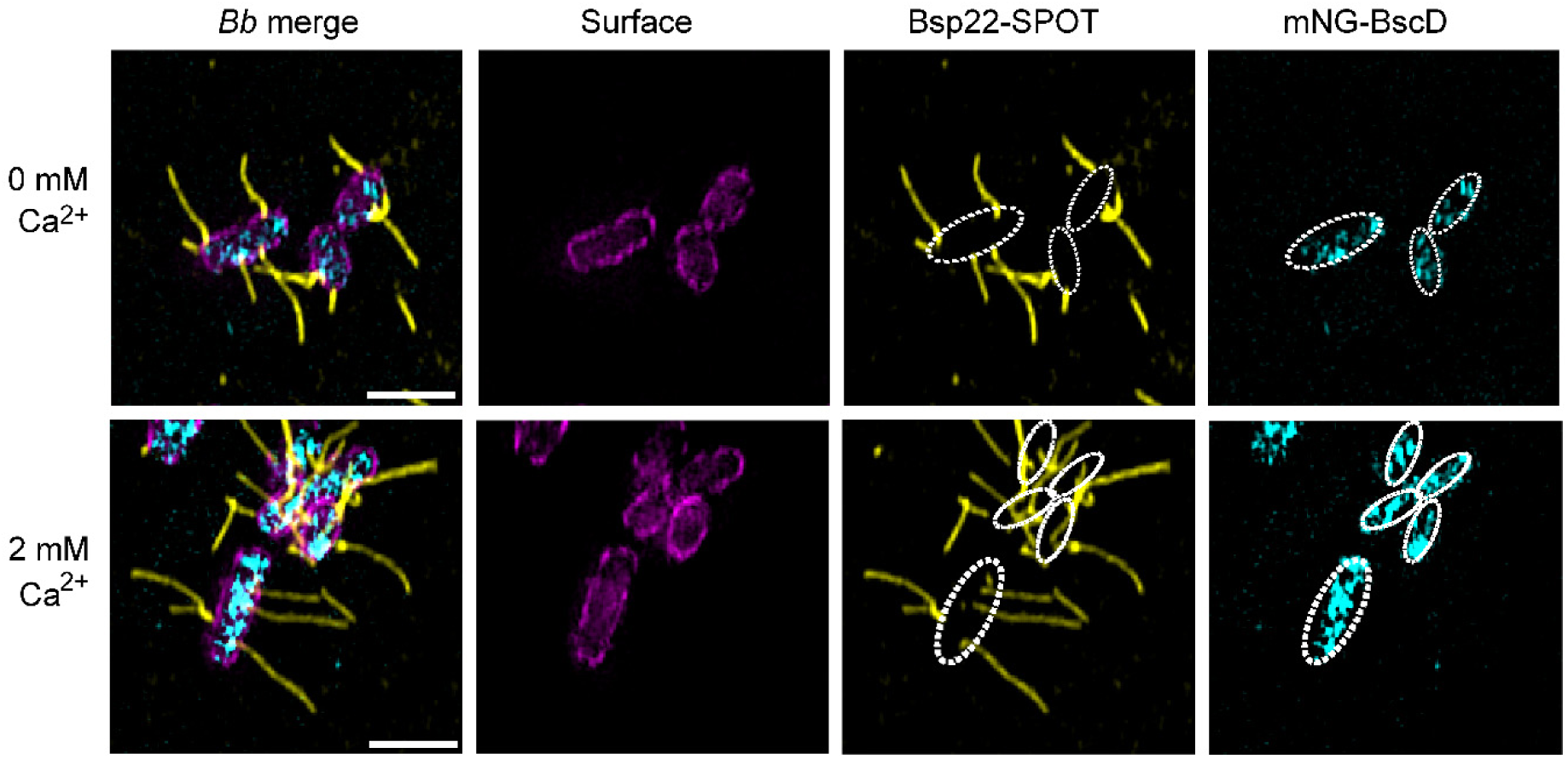
Bsp22 filaments are less abundant than injectisome rings, and their presence is not influenced by calcium regulation. *Bb bsp22*^SPOT^/ mNG-*bscD* cells were cultured on coverslips for 3 h in the absence or presence of 2 mM Ca^2+^ ions. Bsp22 (yellow) was visualized with Spot-label ATTO594, while the *Bb* outer surface (magenta) was stained with a rabbit anti-*Bordetella* serum followed by anti-rabbit IgG-DyLight 405 conjugate. BscD (cyan) was detected as mNeonGreen (mNG) fluorescence. SIM images of a single focal plane are representative of 3 independent experiments. Scale bars, 2 μm.

### Bsp22 filaments interact with the host cell surface and align themselves parallel to the host cell cilia

In the next attempt to comprehend the regulatory mechanisms that govern the formation of Bsp22 filaments, we decided to visualize them during *Bordetella* infection of host cells. For this purpose, we prepared a non-toxic derivative of *Bb bsp22*^SPOT^, *Bb bsp22*^SPOT^ / Δ*bteA*, by deleting the open reading frame for BteA, and verified that this mutation did not disrupt the formation of Bsp22 filaments (Figure S4). Using this strain, we infected HeLa cells expressing a plasma membrane-targeted blue fluorescent protein mTagBFP2, HeLa-PM-BFP, so that we could track their surface. Three hours post-infection, samples were fixed and stained for Bsp22 protein and bacteria. We observed a relatively low number of Bsp22 filaments per bacterial cell (Figure 4A), which were shorter than those observed on glass coverslips. The Bsp22 filaments exhibited different orientations, extending upwards, away from, and towards the HeLa-PM-BFP cell surface. Interestingly, when oriented toward the HeLa cell plasma membrane, they often formed clusters of two or three filaments that closely interacted with the host cell surface, as illustrated in Figure 4B. This interaction resulted in the formation of short physical bridges between the bacterium and the host cell surface.

**Figure 4.**
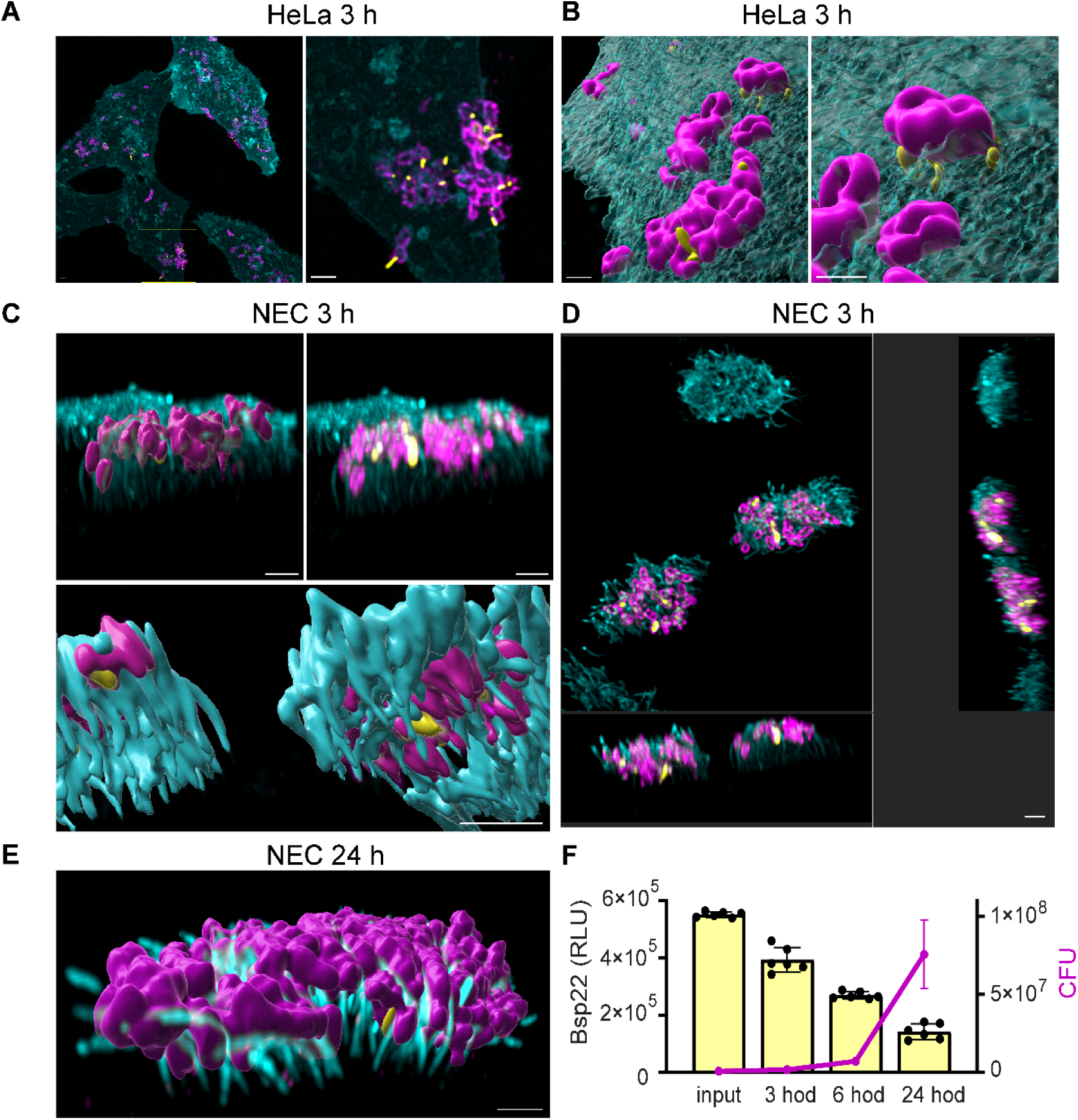
Bsp22 filaments interact with the host cell surface and align themselves parallel to host cell cilia. (A) Bsp22 filaments point in various directions during the infection of non-ciliated cells. HeLa-PM-BFP cells were infected by Bb bsp22SPOT/ ΔbteA for 3 h. Bsp22 (yellow) was visualized with Spot-label ATTO488, while the Bb outer surface (magenta) was stained with a rabbit anti-Bordetella serum followed by anti-rabbit IgG-AF647 conjugate. HeLa-PM-BFP cell surface (cyan) was detected as mTag-BFP2 fluorescence. Maximal intensity projections of confocal images in Z (Z-max) are presented and are representative of 3 independent experiments. Scale bars, 2 μm. (B) A 3D-surface rendering of the portion of the image shown in (A) performed in Imaris illustrates the interaction of Bsp22 filaments with the surface of the HeLa-PM-BFP. Scale bars, 2 μm. (C) Bsp22 filaments are oriented parallel to the host cell cilia during infection of the differentiated nasal epithelia. The apical surfaces of nasal epithelia were infected by *Bb bsp22*^SPOT^/ *ΔbteA* for 3h. Bsp22 (yellow) and *Bb* cell surface (magenta) were stained as described in Figure 3 legend. Cilia (cyan) were stained with an anti-acetylated tubulin antibody followed by anti-mouse IgG-AF488 conjugate. 3D-rendering of all channels of confocal images and their surfaces was performed in Imaris. Images are representatives of three independent experiments. The bottom panel image shows the bottom side of the cilia. Scale bars, 2 μm. (D) Orthogonal view of the same image as in (C) to demonstrate alignment of Bsp22 filaments with cilia. Scale bar, 2 μm. (E) *Bb bsp22*^SPOT^/ *ΔbteA* cells multiply during infection and fill in the space between cilia. 3D-rendering of all channels and their surfaces is representative of three independent experiments. Bsp22 (yellow) and *Bb* cell surface (magenta) were stained as described in Figure 3 legend, cilia (cyan) were stained as stated above in (C). Scale bar, 2 μm. (F) Bsp22 protein production is down-regulated over the course of infection. The apical surfaces of nasal epithelia were infected by *Bb bsp22*^HiBit^/ Δ*bteA* for the specified duration. Luminescence measurements were employed to assess the quantities of Bsp22 protein, while bacterial CFU were determined through the plating method. Shown are mean values ± SD from three biological replicates, each with two technical replicates.

To visualize Bsp22 filaments in the context of the airway ciliated epithelium, we utilized air-liquid interface (ALI) culture system of the human nasal epithelium [29, 30]. Primary nasal cells from healthy donors were cultured under ALI conditions for four weeks to allow the formation of a well-differentiated epithelium prior to infection with *Bb bsp22*^SPOT^ / Δ*bteA.* Differentiated epithelium was then infected for 3, 6, and 24 hours and stained to identify cilia with an anti-acetylated-α-tubulin, bacteria, and Bsp22 filaments. Consistent with previous findings [31] and as shown in Figure 4C-E, the bacteria specifically colonized host cell cilia. At 3 hours post-infection, *Bb bsp22*^SPOT^ / Δ*bteA* cells attached to the tips of the cilia, while the surrounding non-ciliated cells remained free of bacteria. Over the course of the experiment, bacteria multiplied and grew in the cilia clusters, gradually advancing to the base of the cilia (Figure S5 and Figure 4E). By 24 hours post-infection, the space between the cilia was entirely occupied by bacteria (Figure 4E). When Bsp22 filaments were visualized, their number was low, and they were aligned parallel to the host cell cilia (Figure 4C-E and Figure S5), directed towards the host cell plasma membrane. However, assessing filament interaction with the cell body beneath the cilia was challenging due to sample thickness. Interestingly, a decline in the number of Bsp22 filaments was evident in microscopy images as infection progressed. To assess this decrease over the course of infection, we created a reporter strain of *Bb* RB50, *Bb bsp22*^HiBit^ / Δ*bteA*, carrying a HiBit-tag inserted between amino acids M138 and A139 at the previously identified Bsp22 permissive site. A split-luciferase system was then employed for quantifications of Bsp22, utilizing the high-affinity functional complementation between a small 11-amino acid HiBit-tag and the larger 18-kDa LgBit fragment [32]. Consequently, luciferase activity was measured during the infection in *Bb bsp22*^HiBit^ / Δ*bteA* -infected epithelium after the addition of LgBit and substrate. As shown in Fig. 4F, the levels of luciferase activity in samples declined throughout the infection, while the number of bacteria recovered from the same samples increased. This observation confirms a decrease in the production of Bsp22 over the course of infection.

Taken together, the interaction with host cells constrained the growth of Bsp22 filaments, leading to the formation of shorter Bsp22 filaments. During the infection of non-ciliated cells, some of these filaments closely interacted with the host cell surface, aligning with the function of Bsp22 as a T3SS needle tip complex. In the context of nasal epithelial cell infection, *B. bronchiseptica* multiplied in the cilia clusters and Bsp22 filaments aligned parallel to the cell cilia, with their numbers decreasing as the infection progressed.

## Discussion

In this study, we investigated the formation and dynamics of the T3SS tip complex formed by the Bsp22 protein in *B. bronchiseptica* and *B. pertussis*, utilizing super-resolution fluorescence imaging with fluorophore-labeled nanobodies and biarsenic probes. Considering the distinct categorization of Bsp22 within the T3SS tip complex families [21], such exploration is crucial for understanding the assembly and function of diverse T3SS injectisome families. Our examination of bacteria grown on abiotic surface revealed that the Bsp22 protein forms continuously elongating, and flexible filaments extending up to several µm from bacterial cells. During infection of host cells, we observed the adaptive behavior of Bsp22 filaments. Interaction with non-ciliated cells resulted in the formation of short physical bridges between the bacterium and the host cell surface, while colonization of ciliated cells resulted into the alignment of Bsp22 filaments parallel to the cell cilia. Interestingly, a reduction in the number of Bsp22 filaments was observed as the infection of nasal epithelium progressed, hinting at a potential role for these filaments in the early stages of infection and suggesting their regulation by the host environment.

The T3SS injectisome in classical bordetellae distinguishes itself from those of other bacteria due to its elongated extracellular component. This characteristic is reminiscent of T3SS injectisomes found in phytopathogens, which, in contrast to forming short needle filaments, create flexible pilus-like structures known as the Hrp pilus, reaching lengths of several µm [33, 34]. Additionally, T3SS injectisomes of Attaching and Effacing Pathogens (A/E pathogens), such as EPEC (enteropathogenic *E. coli)*, EHEC (enterohemorrhagic *E. coli*), and the murine pathogen *C. rodentium*, also form an extended structure known as the EspA filament [17, 35, 36]. The EspA filament serves to connect the needle filament to the translocon pore in the host cell membrane and is formed through the polymerization of a tip protein, EspA [17, 19, 20, 37, 38].

Despite the lack of significant sequence similarity between tip proteins Bsp22 and EspA, and definition of Bsp22 as founding member of novel T3SS-tip complex family [21], the structure of monomeric Bsp22 predicted by AlfaFold resembles the monomeric EspA [19, 20, 39]. Both proteins exhibit a coiled-coil structure formed between their N- and C-terminal α-helical segments, separated by an extended, ordered loop region. In EspA, this loop region accommodates hypervariable amino acids responsible for antigenic polymorphisms and allows for amino acid insertions [26, 27]. This study demonstrated a similar permissive capability of the corresponding loop in Bsp22, which allowed for peptide display on the surface of Bsp22 filaments and facilitated the detailed super-resolution imaging of Bsp22 filaments using commercially available nanobodies.

We show that *B. pertussis*, similarly to *B. bronchiseptica*, produces long and flexible Bsp22 filaments that protrude up to several µm from bacterial cells. The number of Bsp22 filaments per bacterium is lower compared to the estimated numbers of EspA filaments in EPEC (12 per bacterium) [40], as well as the estimated numbers of injectisomes in *Salmonella* (10-100 per bacterium) [12] or *Yersinia* (30–100 per bacterium) [14]. Additionally, Bsp22 filaments were observed to be less abundant than *Bordetella* injectisome inner ring protein BscD, prompting questions into the regulatory mechanisms initiating *Bordetella* injectisome assembly and Bsp22 filament formation. Nevertheless, once the initiation of Bsp22 filament formation occurs on an abiotic surface, the filament grows continuously, in contrast to needle filament with a defined length control mechanism [41, 42]. This lack of control length mechanism mirrors observations in EspA filaments, where length modulation depends solely on the availability of intracellular EspA subunits [43]. We also demonstrated that newly secreted Bsp22 subunits are added at the distal end of the filament, resembling the mechanisms seen in flagellar [44] and also EspA [43] filament formation, but differing from the basal-growth pattern of type I and type IV pili [45].

During host cell infection, Bsp22 filaments adapted. They displayed controlled growth, likely influenced by interactions with the host cell and/or connection to plasma membrane-inserted translocators BopB [46] and BopD [47]. As only subset of Bsp22 filaments interacted with the cell surface during HeLa cell infection, while others had different orientations, signaling rather than physical prevention were at play. Notably, during infection of nasal epithelium, where bacterial colonization was confined within cell cilia, Bsp22 filaments aligned parallel to the cilia. Although the reason for this orientation is yet to be elucidated, mechanical stress during cilia beating could contribute to this arrangement. As infection in nasal epithelia progressed, the numbers of Bsp22 filaments and the production of Bsp22 protein decreased. This implies that host cell signals may trigger cascades leading to the inhibition of Bsp22 expression. Alternatively, essential signals present in *Bordetella* cultivation media might be lacking. Elevated c-di-GMP levels, known to enhance biofilm formation while suppressing Bsp22 expression [48–50], could potentially serve as one of the signaling mechanisms in colonizing bacteria. These observations also suggest a role for Bsp22 filaments in early stages of infection, paralleling previous reports on the dynamic behavior of EspA filaments by EPEC, speculated to facilitate the traversal of A/E pathogens through the host cell mucus barrier [17, 51].

In summary, our study offers a detailed description of *Bordetella* needle tip filaments behavior on abiotic surfaces and during host cell infection. This was accomplished by incorporating short peptide tags into Bsp22 without compromising injectisome functionality and fluorescence imaging. Through this approach, we investigated the behavior of Bsp22 filaments, highlighting their distinctive features compared to pentameric tip complexes and their similarities with EspA filaments of A/E pathogens. Importantly, we uncovered context-specific and environment-influenced dynamic modulation of Bsp22 filaments, highlighting the adaptability of the *Bordetella* T3SS injectisome.

## Materials and Methods

### EXPERIMENTAL MODEL AND SUBJECT DETAILS

#### Cell lines

HeLa cells (ATCC CCL-2, human cervical adenocarcinoma) and HeLa-PM-BFP (HeLa cells expressing plasma membrane-targeted blue fluorescent protein, mTagBFP2, see below) were cultured in Dulbecco’s Modified Eagle Medium (DMEM) with 10% heat-inactivated fetal bovine serum (FBS, DMEM-10%FBS) at 37°C and 5% CO_2_. For microscopy purposes, DMEM without a phenol-red indicator, supplemented with either 10% or 2% heat-inactivated FBS (DMEM-noPhenolRed-10%FBS and DMEM-noPhenolRed-2%FBS, respectively), to minimize cell autofluorescence was used.

HeLa-PM-BFP stable cell line was generated by lentiviral transduction of the parental HeLa cell line. The pLJM1-PM-BFP vector used for virus production was prepared by subcloning the plasma membrane targeting sequence, Rpre [52] in frame with mTagBFP2 coding sequence at its C-terminus. VSV-pseudotyped viruses were then produced by co-transfecting 6 µg of pLJM1-PM-BFP, 6 µg of pCMV-VSV-G, and 6 µg of psPAX2 plasmids into 293T cells grown in a 10-cm dish using Lipofectamine 2000 (Invitrogen). The cell culture supernatant was collected 48 h after transfection and used to transduce parental HeLa cells in the presence of polybrene (8 µg/mL). Twenty-four hours after transduction, cells were split, selected by puromycin (0.5 µg/mL), and sorted by flow cytometry to obtain single-cell clones of HeLa-PM-BFP cells.

The feeder cell line for nasal epithelial cells, 3T3-J2 (Kerafast EF3003, embryonic mouse fibroblasts), was cultured in DMEM with 10% bovine calf serum (BSC) and antibiotics (0.1 mg/mL streptomycin, 1000 U/mL penicillin). To arrest cell proliferation of feeder 3T3-J2 cells, mitomycin C was added to the medium at a final concentration of 4 μg/mL, and cells were incubated for 2 hours at 37°C and 5% CO_2_.

#### Air-Liquid Interface (ALI) cultures of human nasal epithelial cells

Nasal epithelial cells (NEC) were collected from the anterior nares of healthy donors using a cytology brush. After digestion with TrypLE Select (Gibco) for 5 min at 37°C, the resulting single cells were centrifugated (5 min; 350 g), and gently resuspended in the NEC medium, specified in Table S2. To allow for their conditional reprogramming, NEC cells were seeded into a T25 flask on mitomycin-treated 3T3-J2 fibroblasts in the NEC medium, as previously reported [30]. Once the NEC population had sufficiently expanded, 5 × 10^4^ cells were plated onto the apical side of collagen-coated 6.5-mm Transwell filter (Corning Costar) in 200 µL of apical and 600 µL of basolateral NEC medium without fungin. After 72 h, during which the cells reached confluency, the apical medium was removed (air-lifting), and the basolateral medium was replaced by the differentiation ALI medium (Table S2). The ALI medium was replaced three times per week. The air-lifting defined day 0 of the ALI culture, and the experiments were conducted between days 25 to 28. To prevent mucus accumulation on the apical side, the cells underwent a 30-minute apical wash with phosphate-buffered saline (PBS) every 3 days starting from day 14 of the ALI culture. The differentiation and integrity of ALI cultures were verified through visual inspection and measurement of transepithelial electrical resistance (TEER) before each experiment. Additionally, on day 23 of the ALI culture, the basolateral ALI medium was replaced with the ALI medium without antibiotics, with a minimum of 2 medium changes occurring prior to the experiment.

#### Bacterial strains

The bacterial strains used in this study are detailed in Table S3. Plasmid construction was carried out using *E. coli* strain XL1-Blue, while plasmid transfer into *B. bronchiseptica* RB50 or *B. pertussis* B1917 was performed by bacterial conjugation using *E. coli* strain SM10λpir. *E. coli* strains were cultivated at 37°C on LB agar or in LB broth, supplemented with 100 µg/mL of ampicillin. Wild-type *B. bronchiseptica* RB50 (*Bb*) and *B. pertussis* B1917 (*Bp*), along with their derivatives, were cultivated on Bordet-Gengou (BG) agar medium (Difco) supplemented with 1% glycerol and 15% defibrinated sheep blood (LabMediaServis) at 37°C and 5% CO_2_ for 48 h (*Bb*) or 72h (*Bp*). For the growth of liquid *Bb* and *Bp* cultures at 37°C, modified Stainer-Scholte (SSM) medium, *Bb*-SSM and *Bp*-SSM, respectively, with reduced concentrations of L-glutamate (monosodium salt) and no FeSO_4_.7H_2_O added, as specified in Table S2, was used. Bacteria in the exponential growth phase, with OD_600_ ∼ 1.5 for *Bb* and OD_600_ ∼ 1.0 for *Bp*, were used for the experiments.

## METHOD DETAILS

### Mutagenesis

Plasmids and oligonucleotides used in this study are listed in Table S4 and Table S5, respectively. Plasmids were constructed using the Gibson assembly strategy [53]. PCR amplifications were performed using Herculase II Phusion DNA polymerase (Agilent) from the chromosomal DNA of *B. bronchiseptica* RB50 or *B. pertussis* B1917 as the template. The coding sequence of mTagBFP2 was amplified from mTagBFP2-pBAD (Addgene, [54]). All constructs were verified by DNA sequencing (Eurofins Genomics). Mutant *B. bronchiseptica* and *B. pertussis* strains were constructed by homologous recombination using the suicide allelic exchange vector pSS4245, as previously described [55]. The presence of the introduced mutations was confirmed by PCR amplification of the relevant regions of the *Bordetella* chromosome, followed by agarose gel analysis and DNA sequencing (Eurofins Genomics).

### Determination of cellular cytotoxicity

HeLa cells were seeded at density 5 x 10^4^ per well in a 96-well plate in DMEM without phenol red indicator and supplemented with 2% FBS (DMEM-noPhenolRed-2%FBS). The next day, HeLa cells were infected by exponential *Bb* culture at the indicated multiplicity of infection (MOI). After centrifugation (5 min; 300 g) to increase infection efficiency, HeLa cells were incubated for 3 hours at 37°C and 5% CO_2_. Cellular cytotoxicity was measured as lactate dehydrogenase (LDH) release into the cell culture medium using the CytoTox 96 assay (Promega), according to the manufacturer’s instructions.

### Preparation of glass slides for fluorescence imaging

High-precision coverslips (#1.5 H, 18×18 mm, Marienfeld) were immersed in 1 M HCl for 1 hour, followed by five rinses in deionized water (dH_2_O) and air-drying. Subsequently, the coverslips were stored in 96% ethanol. When needed, coverslips were coated with 0.01% poly-L-lysine (Boster Biological Technology) for 15 min, washed twice with dH_2_O, and air-dried.

### Visualization of Bsp22^SPOT^ on abiotic surfaces of a coverslip

To balance bacterial growth, different colony-forming units (CFUs) of *Bb* or *Bp* in the *Bb*-SSM and *Bp*-SSM medium, respectively, were applied on each coverslip within a 6-well plate. The CFU amounts used were 1.5 x 10^9^ CFU for a 0.5-hour incubation, 6 x 10^8^ CFU for a 1- and 2-hour incubation, 1.5 x 10^8^ CFU for a 3-hour incubation, and 3 x 10^7^ CFU for a 6-hour incubation. After seeding, the bacteria were centrifuged (3 min; 300 g) to facilitate adhesion and then cultivated at 37°C and 5% CO_2_. At the end of the incubation period, the coverslips were rinsed with PBS and adherent cells were fixed with 4% paraformaldehyde (PFA) for 20 minutes at room temperature (RT). Coverslips were then washed with PBS (3 x 5 min) and blocked with 4% BSA in PBS (4%BSA-PBS) for 1 hour at RT. The SPOT-tag nanobody conjugated with ATTO488 (Spot-label ATTO488) or ATTO594 (Spot-label ATTO594) at a dilution of 1:4,000 and a rabbit anti-*B. pertussis* serum (provided by Dr. Vecerek, Institute of Microbiology, Prague, Czech Republic) at a dilution of 1:1,000 in 1% BSA in PBS (1%BSA-PBS) were applied overnight at 4 °C. The following day, the coverslips were washed in PBS (3 x 5 min) and incubated with anti-rabbit IgG-DyLight 405 conjugate at a dilution of 1:600 in 1% BSA-PBS for 1 hour at RT. After washing in PBS (3x 5 min) and brief rinses in dH_2_O (10x), coverslips were mounted on glass slides using the Vectashield mounting medium and SecureSeal Imaging Spacers.

### Pulse-chase labeling of Bsp22^TC^

Pulse-chase labeling of *Bb bsp22*^TC^ with the biarsenic reagents ReAsH (ReAsH-EDT_2_) and FlAsH (FlAsH-EDT_2_) was carried out in reducing *Bordetella* labeling medium (BLM) containing 0.5 mM TCEP (tris(2-carboxyethyl)phosphine hydrochloride). BLM, developed in this study, is based on M9 minimal and SSM media, and its composition is detailed in Table S2. *Bb bsp22*^TC^ cells (1.5 x 10^9^ CFU) were pelleted (10 min; 1,200 g), resuspended in BLM, and incubated for 30 min at 37°C to allow reduction of extracellular cysteines. ReAsH was then added at a final concentration of 1 μM, and the bacteria were incubated stationary in an Eppendorf tube at 37 °C. Progress of labeling was checked by wide-field microscopy when 5-μL samples of the culture were spotted onto a coverslip and covered with an agarose pad prepared in PBS. The coverslip was attached to a support and observed under a wide-field microscope (Olympus). When Bsp22 filaments were well-discernible on the cells (after 220 min), ReAsH-EDT_2_ was removed by centrifugation (10 min; 1,200 g). Pelleted bacteria were resuspended in BLM and incubated for 30 min at 37 °C. FlAsH-EDT_2_ was then added at a final concentration of 1 μM, followed by incubation at 37°C. Progress of labeling was again checked by wide-field microscopy of culture spotted onto a coverslip and covered with an agarose pad. When double-labeled Bsp22 filaments were visible (after 60 min), the bacteria were applied onto poly-L-lysine-coated coverslips and allowed to adhere for 10 minutes. After rinsing with PBS once, the cells were fixed with 4% PFA for 20 minutes at RT. The coverslips were washed with PBS (3x 5 min), rinsed in dH_2_O (10x), and mounted using Vectashield onto glass slides.

### Visualization of Bsp22^SPOT^ during HeLa cell infection

The HeLa-PM-BFP cells were seeded at a density of 2.5 x 10^5^ in a 12-well plate using DMEM-noPhenolRed-10%FBS and allowed to grow overnight. The following day, the medium was replaced with DMEM-noPhenolRed-2%FBS, and the cells were infected with *Bb bsp22*^SPOT^ /Δ*bteA* at MOI 50:1 followed by centrifugation (3 min, 300 g) to enhance the infection efficiency. Three hours post-infection at 37°C with 5% CO_2_, the cells were rinsed with PBS and fixed with 4% PFA for 20 minutes at RT. Coverslips were washed with PBS (3 x 5 min), blocked with 4%BSA-PBS for 1 hour at RT, and incubated with Spot-label ATTO488 at a dilution of 1: 5000 and the rabbit anti-*B. pertussis* serum at a dilution of 1:1,000 in 1%BSA-PBS overnight at 4 °C. Subsequently, the coverslips were washed in PBS (3 x 5 min) and incubated with anti-rabbit IgG-AF647 conjugate at a dilution of 1:600 in 1% BSA-PBS for 1 h at RT. Finally, following washing in PBS (3x 5 min) and rinsing in dH_2_O (10x), coverslips were mounted on glass slides using the Vectashield mounting medium.

### Visualization of Bsp22^SPOT^ dynamics in ALI cultures of human nasal epithelial cells

Differentiated ALI cultures of human nasal epithelial cells were apically infected with *Bb bsp22*^SPOT^ /Δ*bteA* using 100 μL of DMEM-noPhenolRed-2%FBS. For 3- and 6-h infections, 5×10^6^ CFU were applied per a Transwell membrane, and for the 24-hour infection, 10^5^ CFU were used. After infection at 37°C with 5% CO_2_, the membranes were rinsed with PBS and fixed with 4% PFA for 20 minutes at 37°C. Following fixation, membranes were washed with PBS (3x 5 min), and samples were permeabilized with 0.2% Triton-X100 (TX-100) in PBS for 10 minutes at RT. Subsequently, membranes were washed with PBS supplemented with 0.05% TX-100 (PBST) and blocked with 4% BSA in PBST for 1 hour at RT. Spot-label ATTO594 at a dilution of 1:5,000, rabbit anti-*B. pertussis* serum at a dilution of 1:500, and mouse monoclonal anti-acetylated tubulin antibody at a dilution of 1:500 in 1%BSA-PBS were applied overnight at 4 °C. The following day, the membranes were washed with PBST (3x 5 min) and incubated with the anti-rabbit IgG-DyLight 405 conjugate at a dilution of 1:600, and the anti-mouse IgG-AF488 conjugate at a dilution of 1:500 in 1% BSA in PBST for 1 hour at RT. This was followed by washes with PBST (3 x 5 min) and brief rinses in dH_2_O (10x). Finally, the membranes were dissected from transwells, placed inside SecureSeal Imaging Spacers attached to the glass slides, covered with the Vectashield mounting medium, and sealed with coverslips.

### Microscopy

Wide-field microscopy was employed to check the progress of pulse-chase labeling of Bsp22^TC^ and to inspect samples obtained with the above-described methods prior to examining them by SIM or confocal microscopy. The Olympus IX83 inverted microscope equipped with the UPLXAPO100XOPH NA 1.45 objective and sCMOS camera Prime 95B (Photometrics, USA) with high quantum efficiency, large field of view, and large pixel size of 11 μm was used. mTagBFP-2 and DyLight405 were observed using the excitation filter 402/15 nm, the emission filter 455/50 nm, and a quadruple dichroic mirror DAPI/FITC/TRITC/CY5. ATTO488 and AF488 were excited through the filter 490/20 nm and captured by the 525/36 nm emission filter. When using ATTO594 and AF568, the excitation filter 572/35 nm and the emission filter 632/60 nm were combined with a triple dichroic mirror DAPI/FITC/TxRed.

Confocal images were acquired with Leica STELLARIS 8 FALCON equipped with wide-range (440 – 790 nm), light laser with pulse picker (WLL PP), and highly-sensitive hybrid detectors operated by the LAS X software. The objective HC PL APO 63x/1.40 OIL CS2; FWD 0.14 MM was used with the type F immersion oil (Leica, 11513859). mTagBFP2 and DyLight405 were excited with the 405 nm cw diode laser and other dyes with the WLL-PLL laser with the following settings: AF488 (491 nm), ATTO488 (501), ATTO594 (602 nm), AF647 (644 nm). Z-stacks were acquired with steps of 0.15 μm, and pixel size was set to meet Nyquist sampling criteria for deconvolution with the Huygens software (SVI, Netherlands).

The DeltaVision OMX imaging platform was used to acquire images with 3D structured illumination (3D-SIM). The system was equipped with the PLAN APO N 60x oil objective, NA 1.42; FWD 0.15; CG 0, four PCO, and an Edge 5.5 sCMOS camera (readout speeds 95 MHz, 286 Mhz, 15-bit, pixel size 6.5 μm). To excite fluorescent proteins and the labels on the antibodies, 405 nm diode and 488 nm OPSL were used in combination with emission filters 435.5/31 and 528/48 nm. SoftWoRx software (Applied Precision, USA) was used for image reconstruction and registration.

### Image processing

Acquired confocal images were deconvolved using Huygens Professional software (SVI). Orthogonal views, 3-D, and surface rendering were prepared in Imaris, the image analysis software (Oxford Instruments). Further image processing, consisting of cropping and brightness/contrast adjustment, was performed in FIJI ([56], NIH Bethesda). Final images were assembled in Adobe Illustrator (Adobe).

### Determination of bacterial colony-forming units and Bsp22 levels in infected ALI cultures of human nasal epithelial cells

Differentiated ALI cultures of human nasal epithelial cells were apically infected with 5×10^6^ CFU of *Bb bsp22*^HiBit^ /Δ*bteA* per Transwell membrane, using 100 μL of DMEM-noPhenolRed-2%FBS, for durations of 3, 6, and 24 hours. To retrieve bacteria for determining colony-forming units (CFU) and Bsp22 protein levels, 100 µL of PBS supplemented with 0.2% TX-100 was added to the Transwell membrane, followed by a 5-minute incubation. The resulting suspension was then pipetted up and down and transferred into a tube, and an additional 100 µL of PBST was applied onto the Transwell membrane. After a 5-minute incubation, the suspension was once again pipetted up and down and transferred to the same tube. The removal of bacteria and epithelial cells from the Transwell membrane was confirmed by wide-field microscopy. For CFU determination, the suspension was plated at various dilutions on BG agar plates, and CFUs were counted after a 3-day incubation period. To assess Bsp22 levels, the Nano-Glo HiBiT Extracellular Detection System (Promega) was utilized following the manufacturer’s instructions. In brief, a 10x diluted suspension was combined with recombinant LgBit protein, furimazine substrate, and Nano-Glo buffer. Luminescence was subsequently measured using the TecanSpark microplate reader (Tecan).

### ETHICS STATEMENT

The sampling of the anterior nares was conducted on healthy donors who provided written informed consent for the use of their cells in research. All studies were carried out in accordance with the principles outlined in the Declaration of Helsinki.

## Acknowledgments

We thank Sarka Knoblochova, MD, and Barbora Pravdova, MSc. for assistance with ALI cultures of NEC, and Tania Romero Allsop. MSc. for preparation of HeLa-PM-BFP cell line. This research was funded by grant 24-11053S of the Czech Science Foundation (www.gacr.cz) and grant Talking microbes - understanding microbial interactions within One Health framework (CZ.02.01.01/00/22_008/0004597) of the Ministry of Education, Youth and Sports of the Czech Republic (www.msmt.cz) to JK. We would also like to acknowledge the support of the project LM2023053 (Czech National Node to the European Infrastructure for Translational Medicine) from Ministry of Education, Youth and Sports of the Czech Republic, and the Light Microscopy Core Facility, IMG, Prague, Czech Republic, particularly the contributions of Ivan Novotny, PhD., supported by MEYS – LM2023050 and RVO – 68378050-KAV-NPUI, for support with the super-resolution and confocal imaging presented herein. The funders had no role in study design, data collection and analysis, decision to publish, or preparation of the manuscript.

## Author contributions

Conceptualization: J.K., I.M.; methodology: I.M.; formal analysis: J.K., I.M..; investigation: I.M.; writing: J.K., I.M.

## Inclusion and diversity statement

Figures were generated using a color-blind accessible palette, promoting inclusion and diversity in data visualization.

## Author Summary

*Bordetella bronchiseptica* and *Bordetella pertussis*, closely related respiratory pathogens that selectively colonize ciliated cells, employ their T3SS injectisome to deliver the BteA effector into host cells. In this study, we visualized the needle tip filament of their T3SS injectisome, a structure formed by the Bsp22 protein. We demonstrate the dynamic extension of the injectisome length, where Bsp22 subunits are added at its distal end. Within nasal epithelium, the resulting flexible Bsp22 filaments align parallel to cilia, showcasing an adaptation to this specific environment. These findings illustrate how *Bordetella* adapts its T3SS injectisome to various environments and host cell interactions during infection.

## Suppl. figures

**Figure S1.**
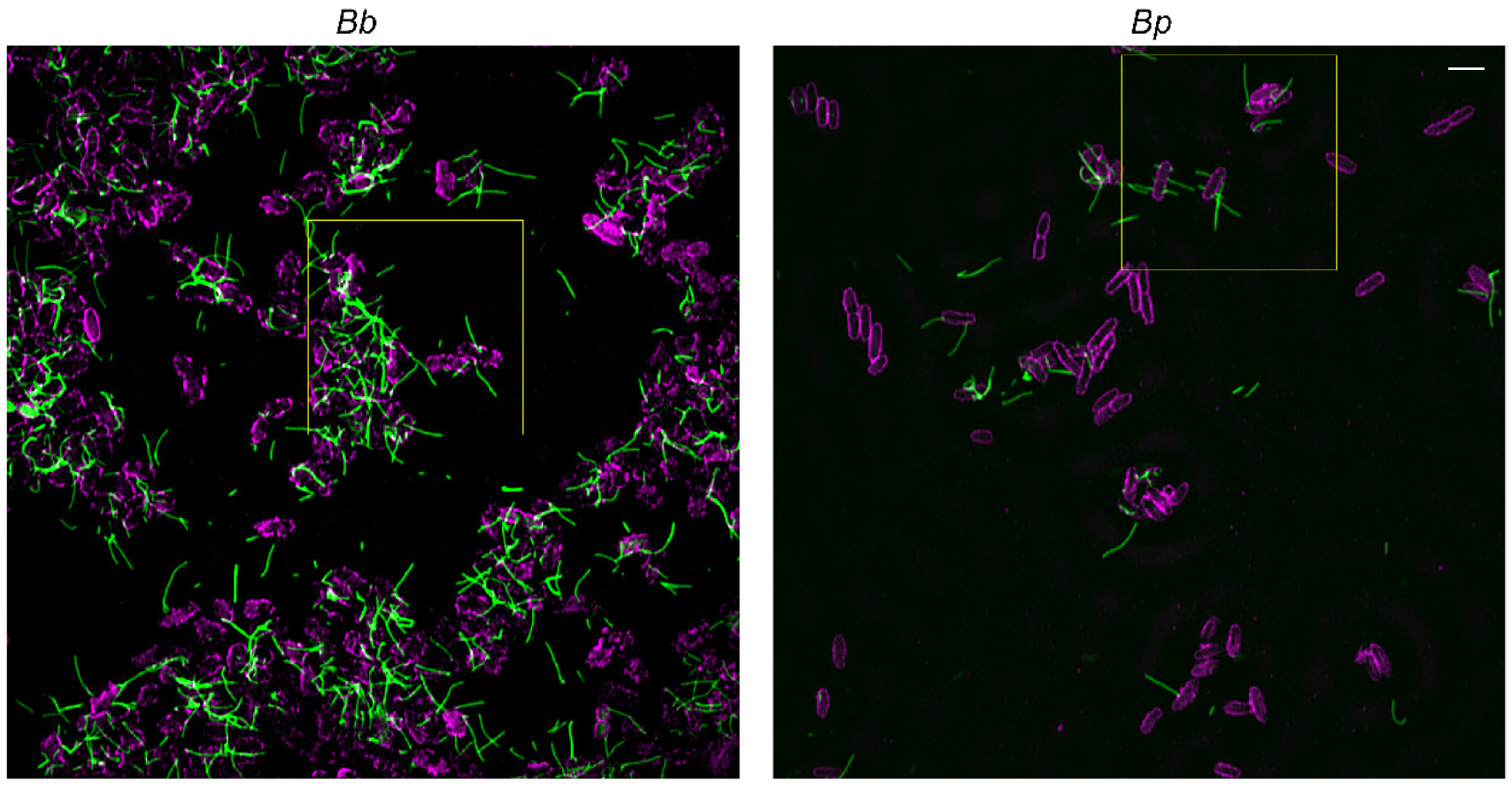
The Bsp22 protein polymerizes into flexible filaments on abiotic surfaces of glass coverslips (related to Figure 1). Cells of *B. bronchiseptica* RB50 (*Bb*) or *B. pertussis* B1917 (*Bp*) carrying *bsp22*^SPOT^ were cultured on coverslips for 3 h. Following fixation, Bsp22 (green) was visualized with the Spot-label ATTO488, while the bacterial outer surface (magenta) was labeled with a rabbit anti-*Bordetella* serum followed by anti-rabbit IgG-DyLight 405 conjugate. The yellow squares highlight the regions magnified in Figure 1B and 1C. The shown SIM images depict a single focal plane and are representative of three independent experiments. Scale bars, 2 μm.

**Figure S2.**
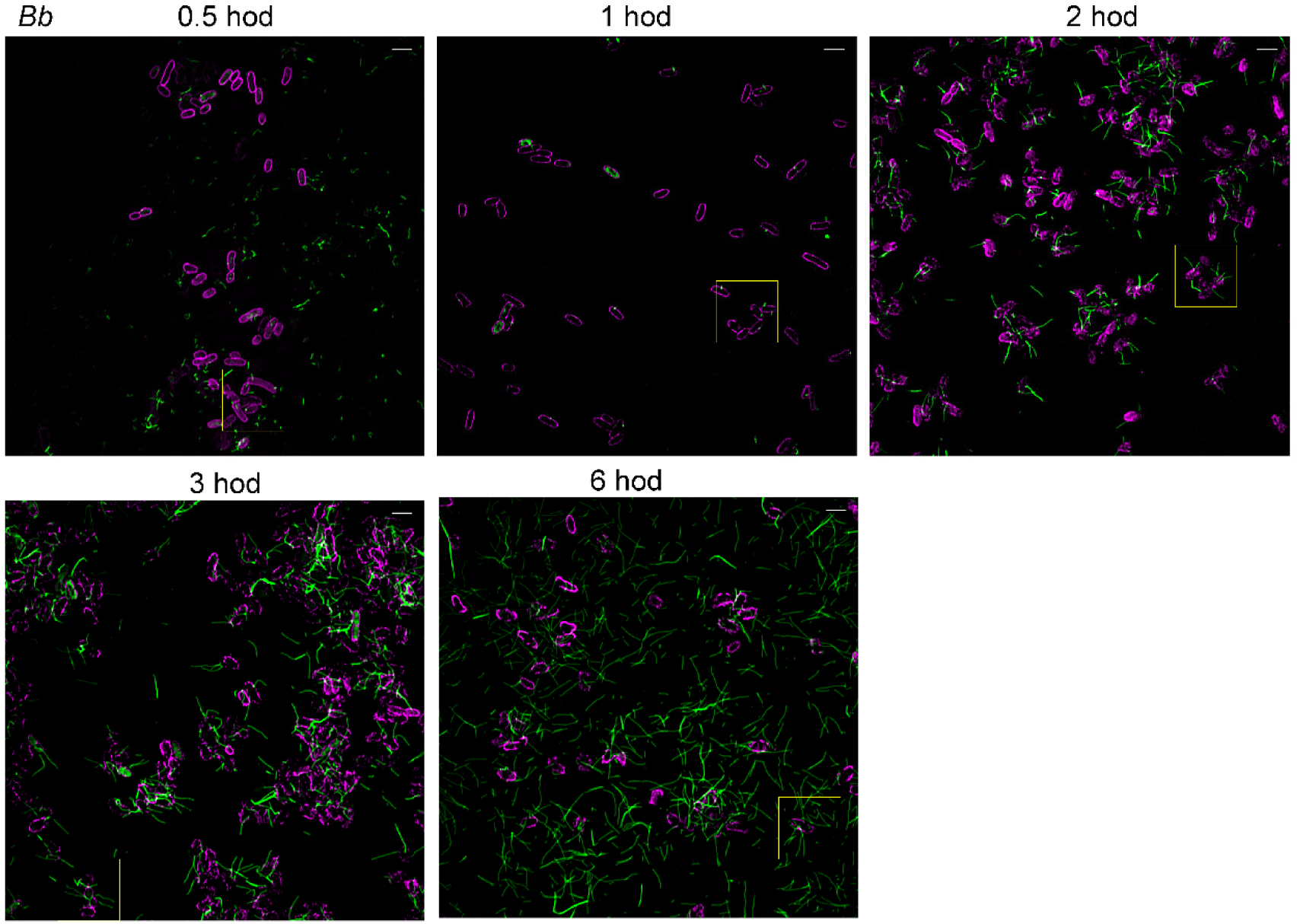
The growth of Bsp22 filaments is continuous (related to Figure 2). Cells of *Bb bsp22*^SPOT^ cultured on coverslips were fixed at the indicated time points. Staining was performed as described in the legend of Figure 1C. Bsp22, green; *Bb* outer surface, magenta. The yellow squares highlight the regions magnified in Figure 2A. The SIM images show a single focal plane and are representative of three independent experiments. Scale bars, 2 μm.

**Figure S3.**
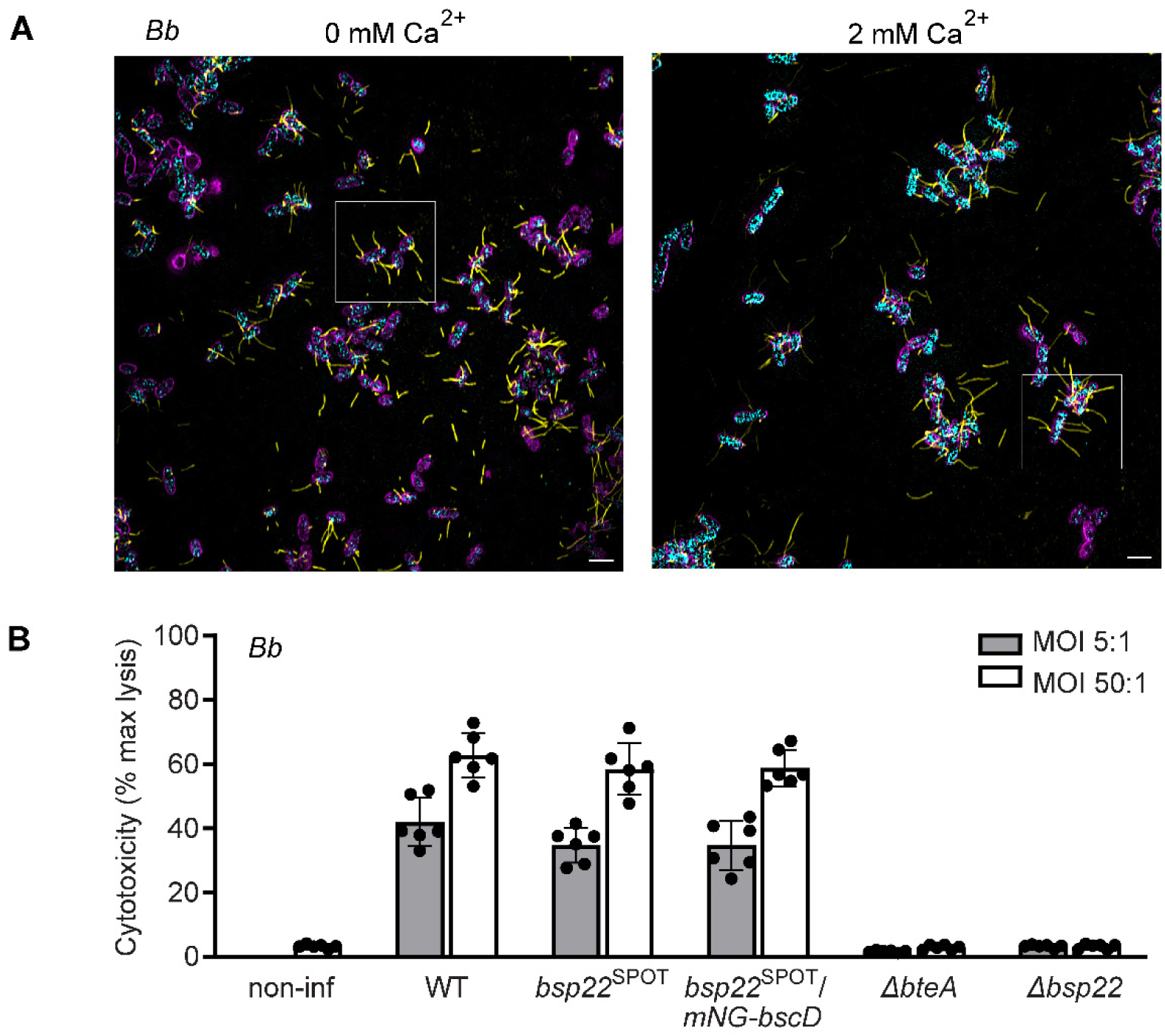
Bsp22 filaments are less abundant than injectisome rings, and calcium regulation does not influence their presence (related to Figure 3). (A) Bb bsp22SPOT/ mNG-bscD cells were fixed on coverslips after cultivation for 3 hours in the absence or presence of 2 mM Ca2+. Bsp22 (yellow) and Bb cell surface (magenta) were stained as described in Figure 3 legend. BscD localization (cyan) was detected as mNG fluorescence. The yellow squares highlight the regions magnified in Figure 3. The SIM images show a single focal plane and are representative of three independent experiments. Scale bars, 2 μm. (B) Tagging of bscD by mNG does not affect the functionality of the Bordetella injectisome. T3SS-dependent cytotoxicity of the indicated Bb strains towards HeLa cells was measured as described in Figure 1B legend. Mean values ± SD from two biological replicates with three technical replicates each are shown.

**Figure S4.**
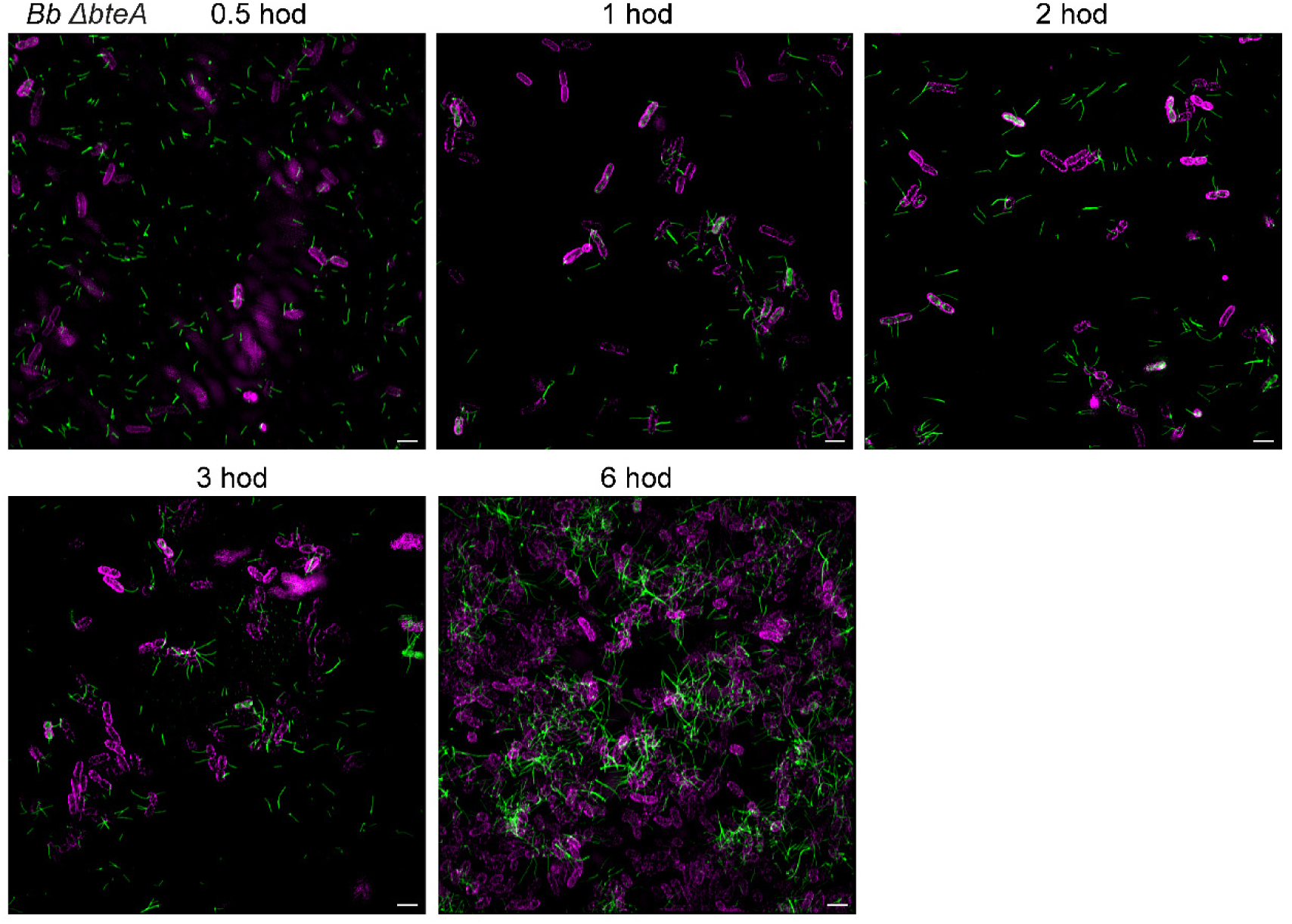
Inactivation of the effector protein BteA does not hinder the formation of Bsp22 filaments. *Bb bsp22*^SPOT^/ Δ*bteA* cells were cultivated in *Bb*-SSM on coverslips and fixed at the indicated time points. Staining was performed as described in Figure 1C legend. Bsp22, green; *Bb* outer surface, magenta. SIM images of a single focal plane are representatives of 3 independent experiments. Scale bars, 2 μm.

**Figure S5.**
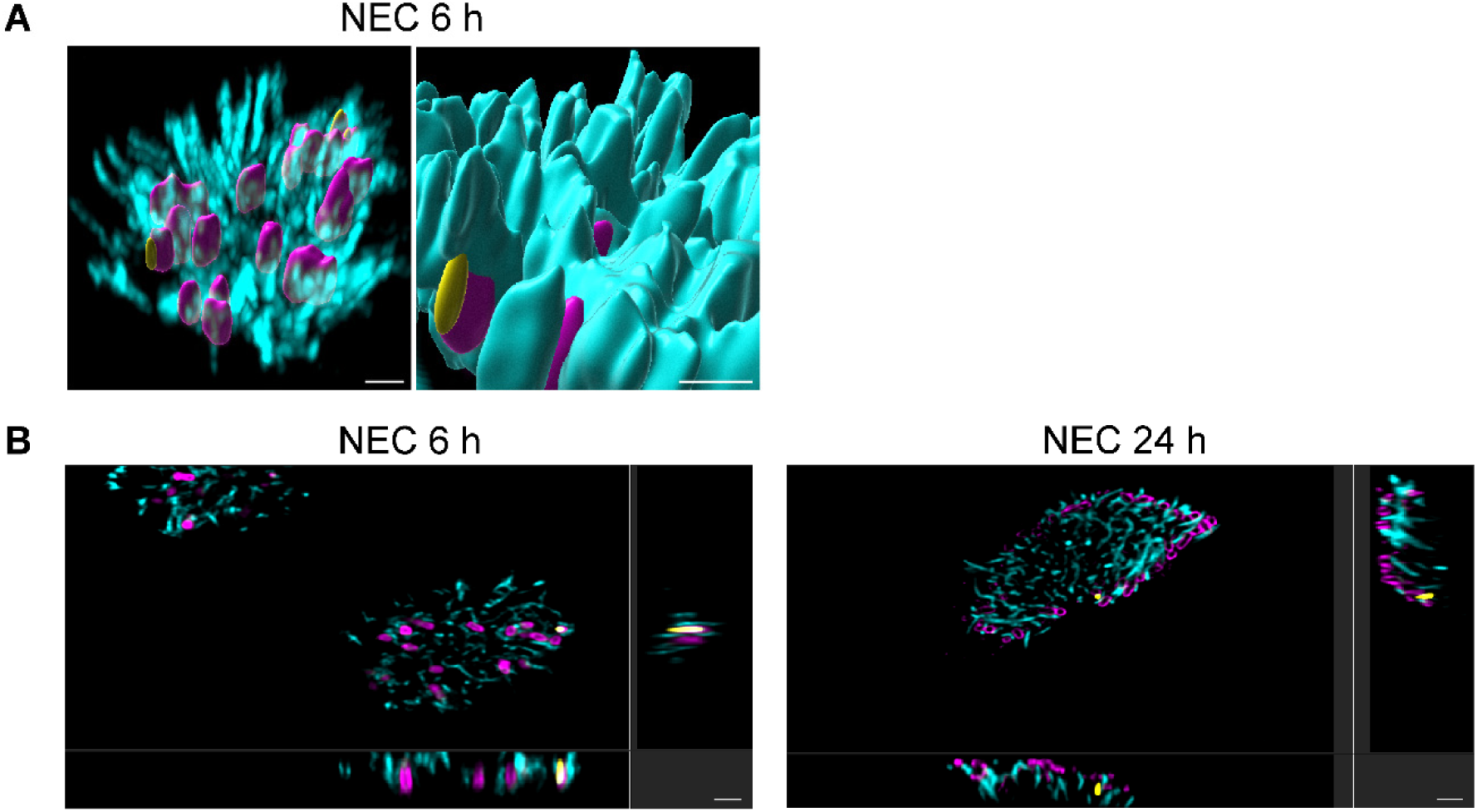
Distribution of Bsp22 filaments and their alignment with cilia at 6- and 24-h intervals following nasal epithelial infection with Bb bsp22SPOT/ ΔbteA. (A) Distribution of Bsp22 filaments in nasal epithelia 6 h post-infection. Bsp22 (yellow) and *Bb* cell surface (magenta) were stained as described in Figure 3 legend. Cilia (cyan) were stained with an anti-acetylated tubulin antibody followed by anti-mouse IgG-AF488 conjugate. 3D-rendering of all channels and their surfaces is representative of three independent experiments. Scale bars, 2 μm. (B) Orthogonal views of infected epithelia 6 and 24 h post-infection. Bsp22 (yellow) and *Bb* cell surface (magenta) were stained as described in Figure 3 legend, cilia (cyan) were stained as stated above in (A). Orthogonal views are representative of three independent experiments. Scale bars, 2 μm.

## Suppl. tables

**S1. Table.**
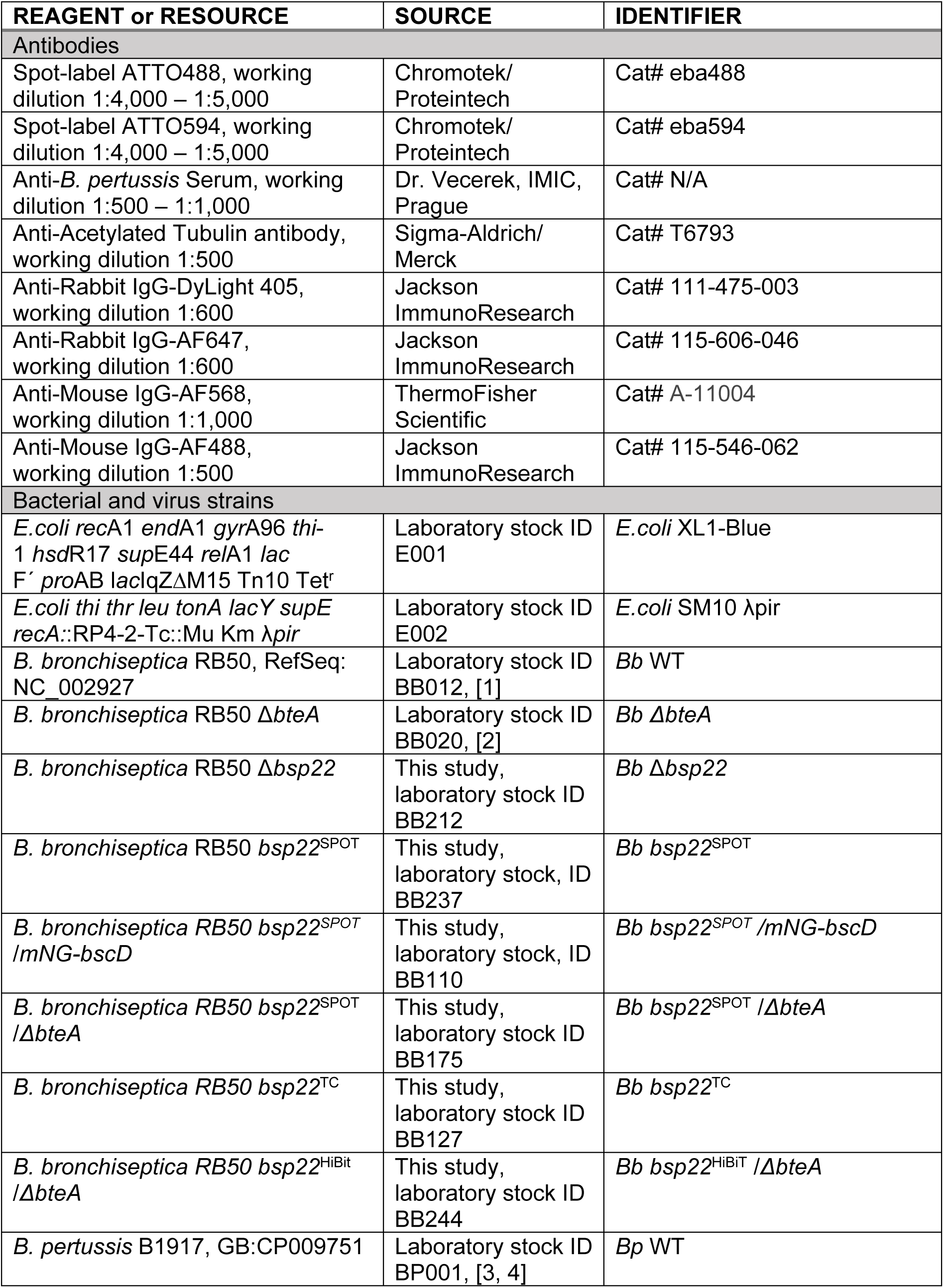
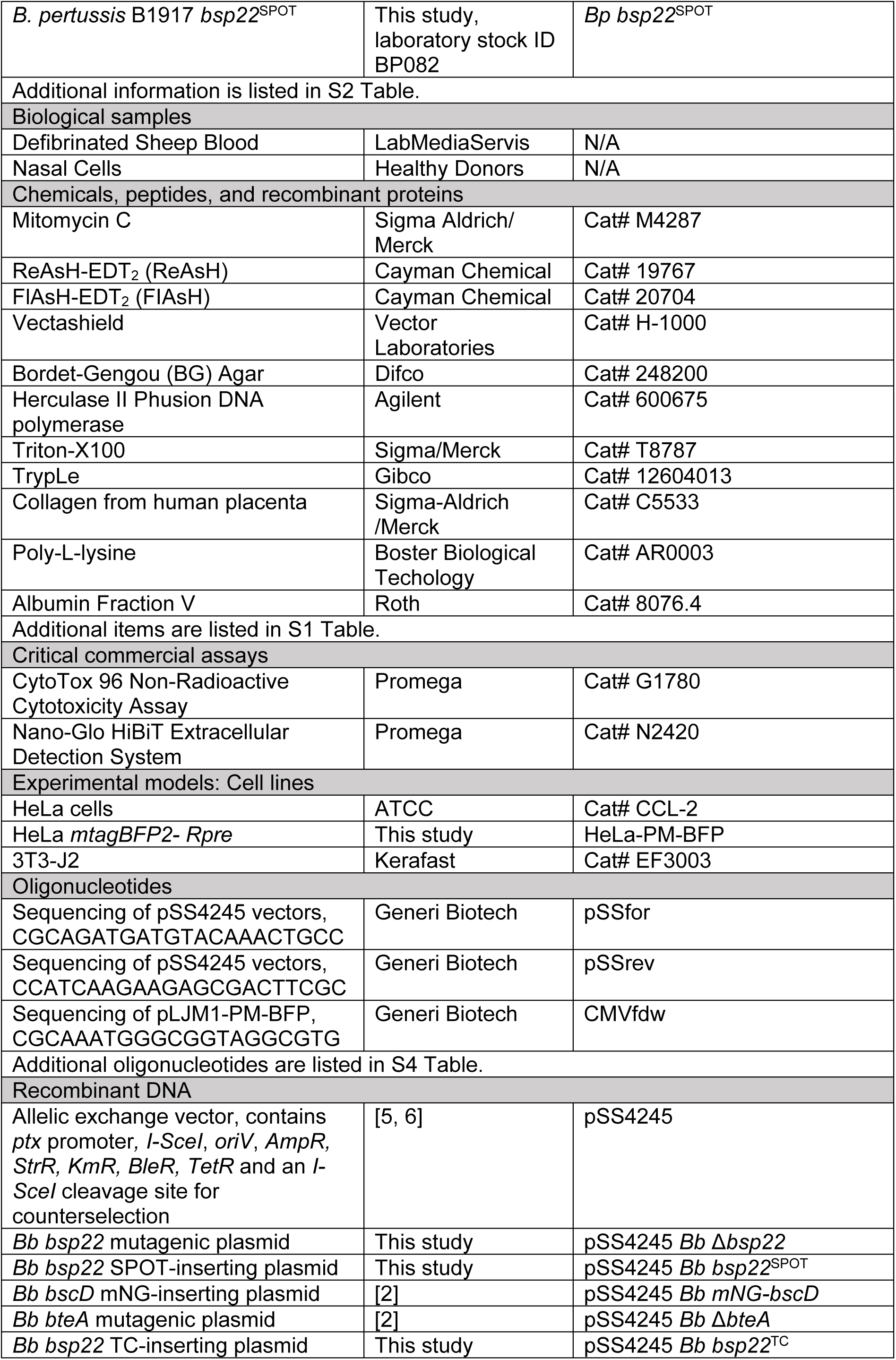
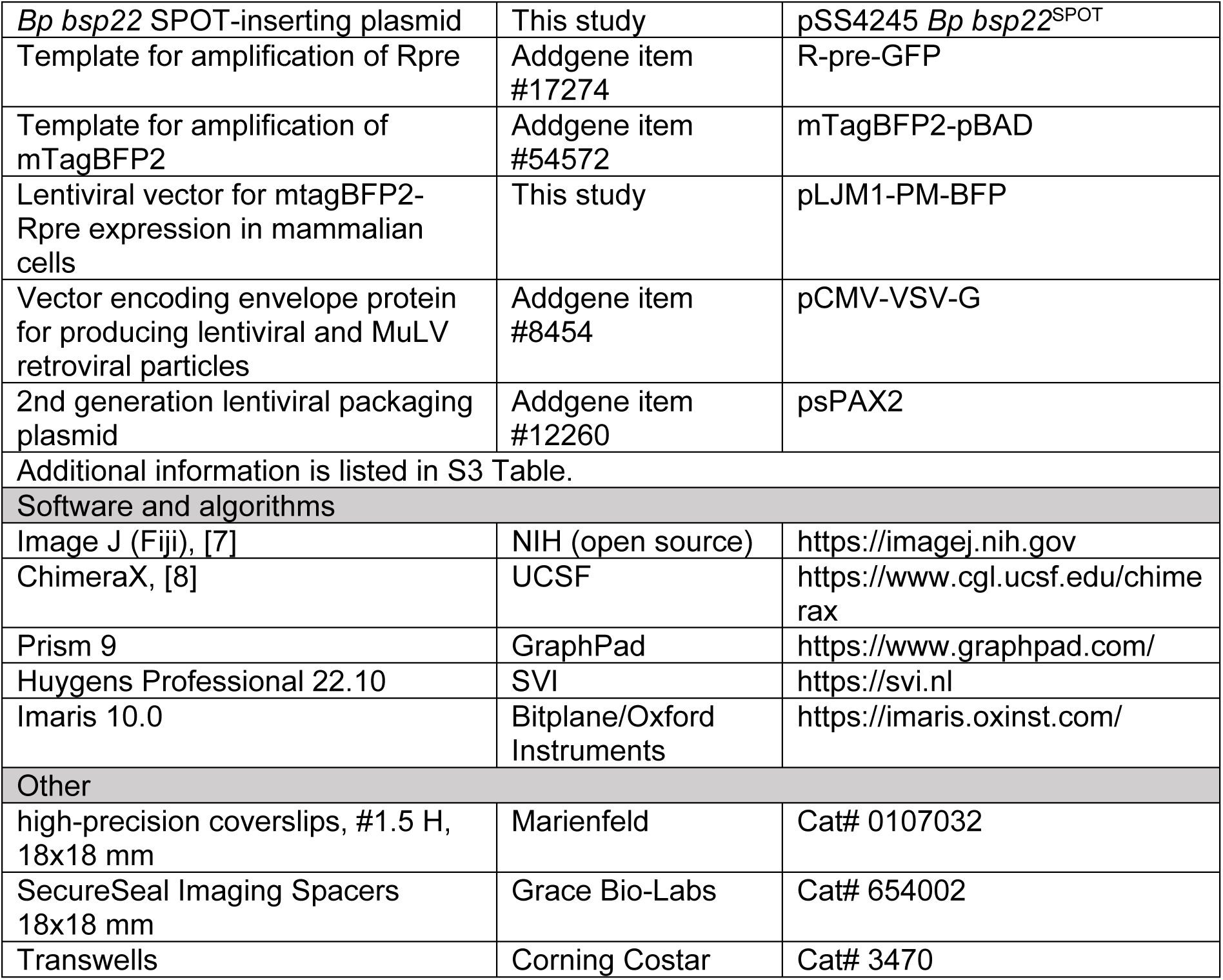
KEY RESOURCES TABLE.

**S2. Table.**
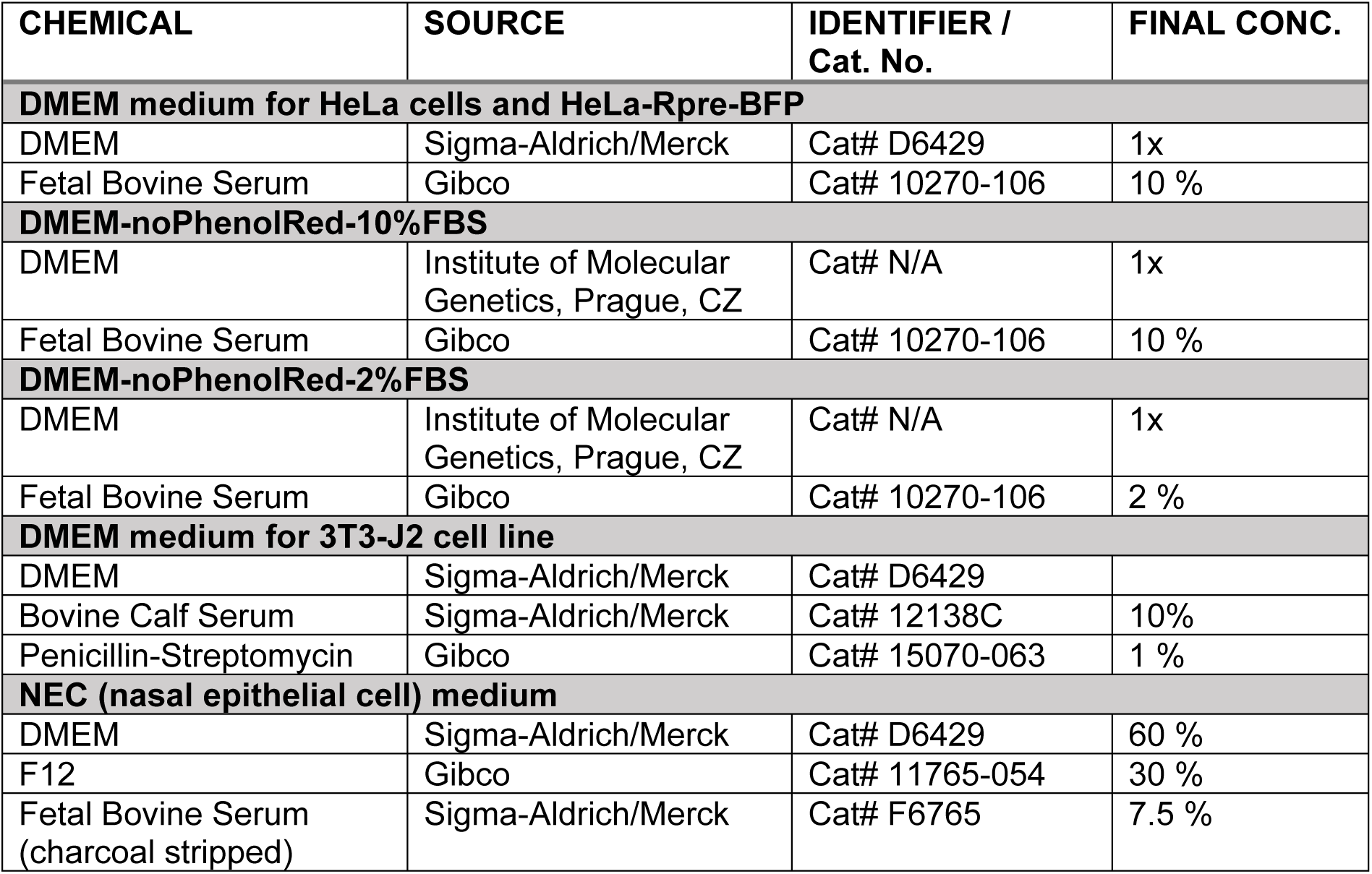
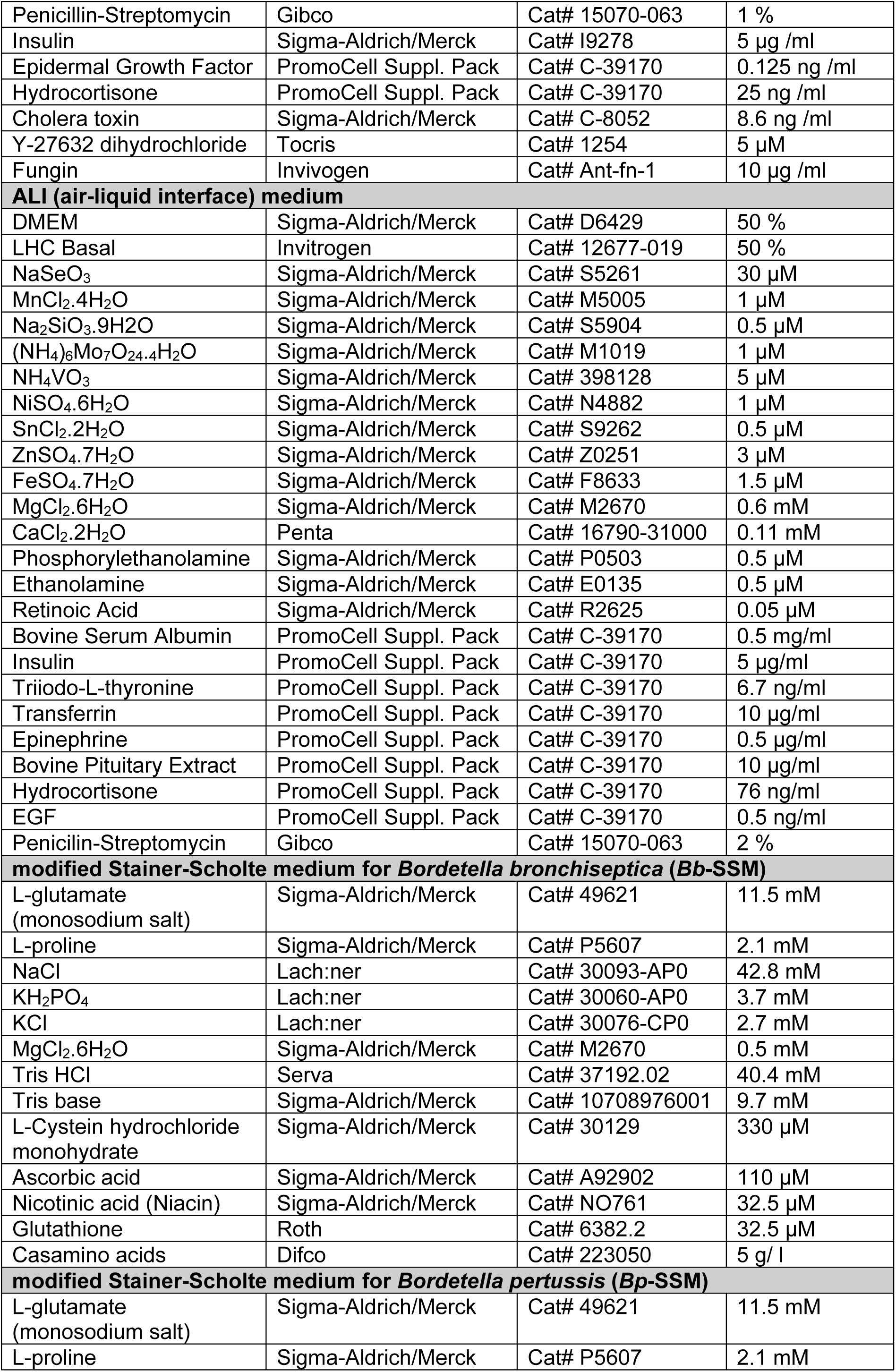
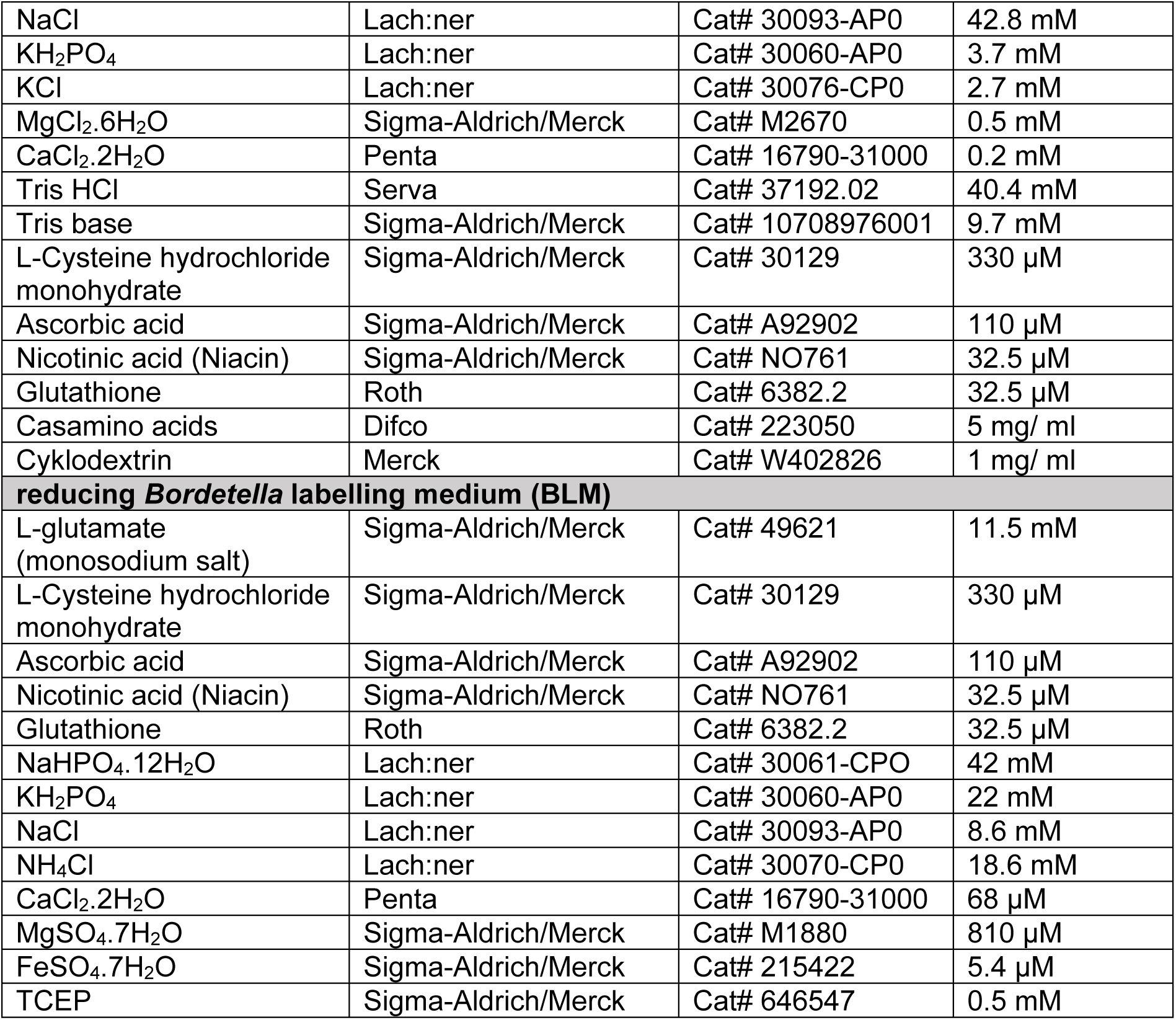
COMPOSITION OF CELL CULTURE AND BACTERIAL CULTIVATION MEDIA.

**S3. Table.**
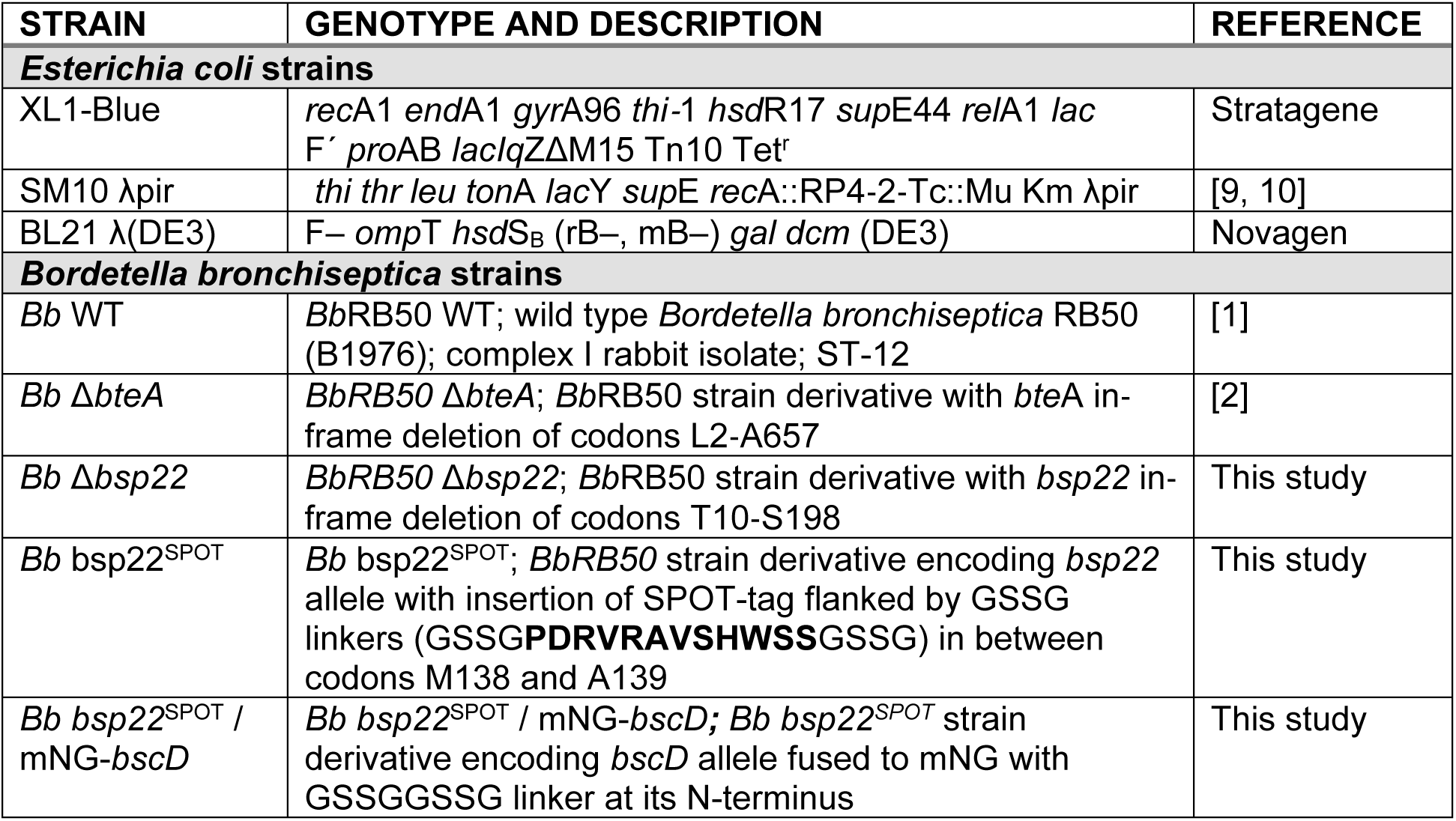
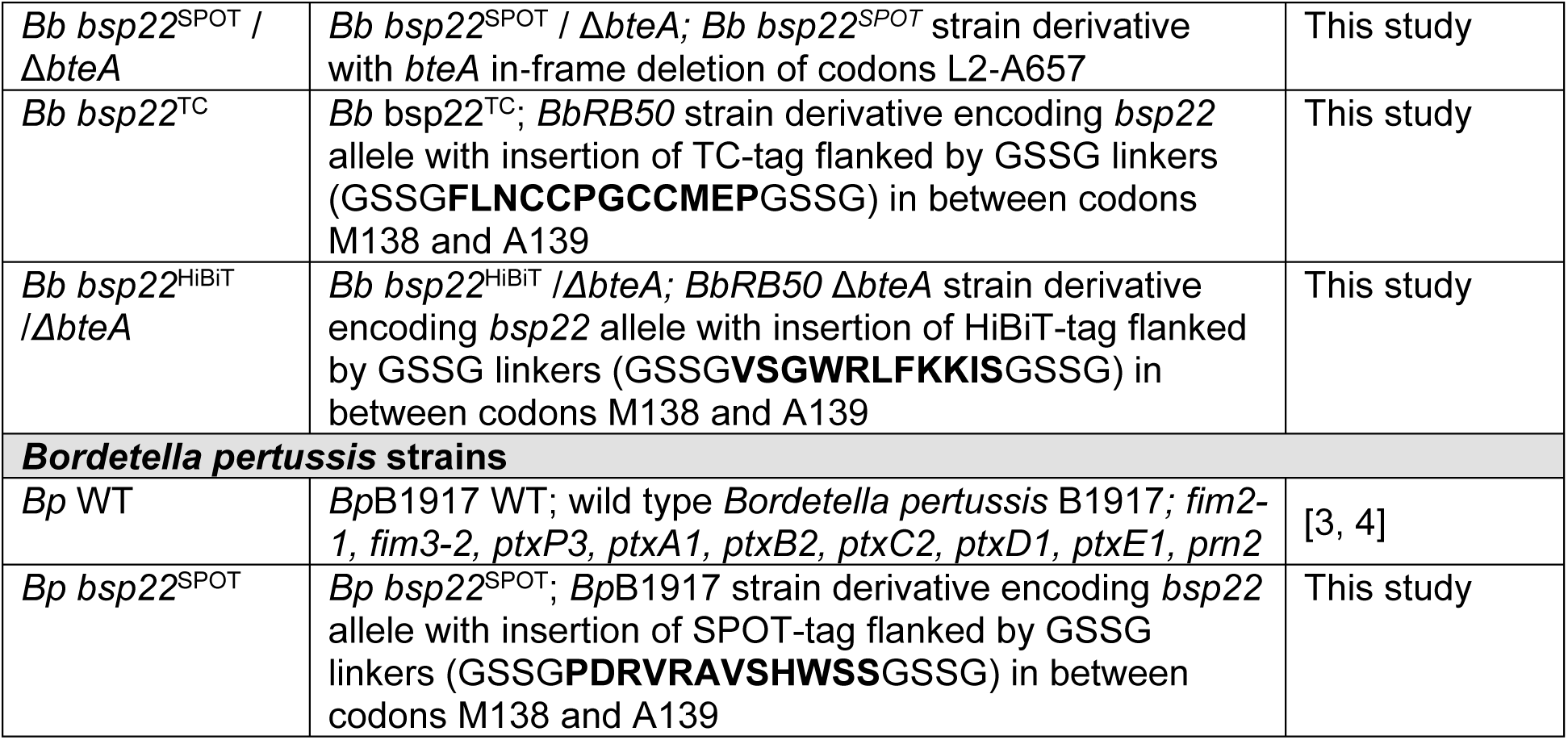
DESCRIPTION OF BACTERIAL STRAINS USED IN THIS STUDY.

**S4. Table.**
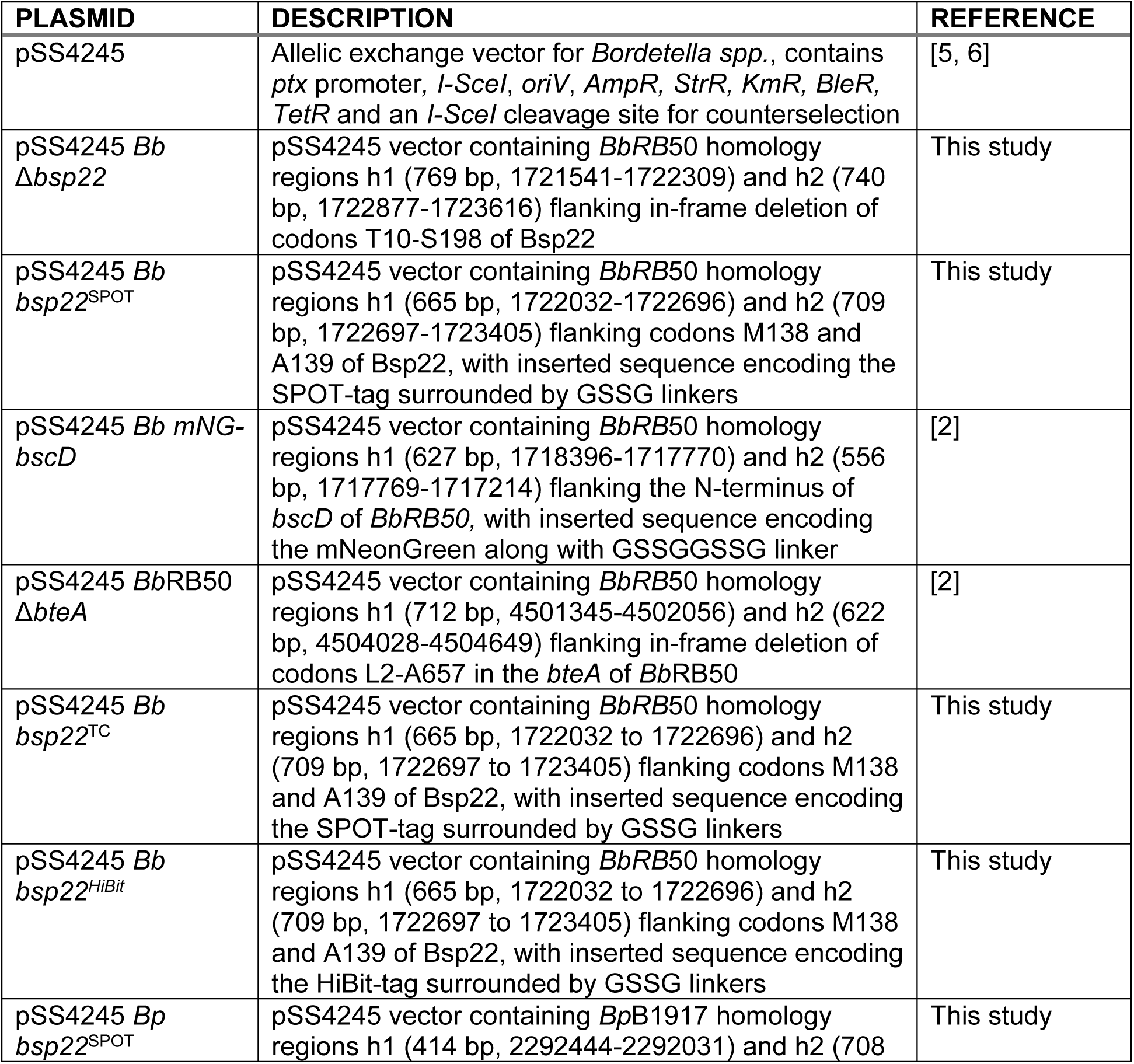
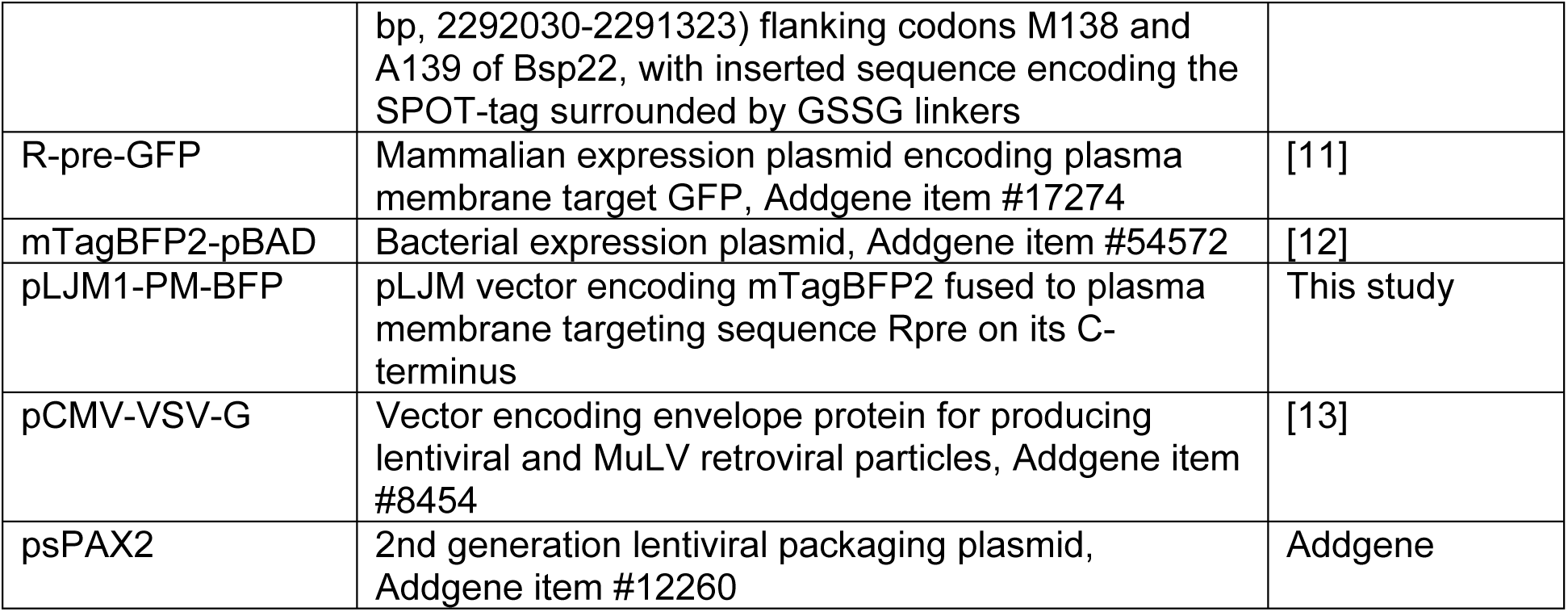
DESCRIPTION OF PLASMIDS USED IN THIS STUDY.

**S5. Table.**
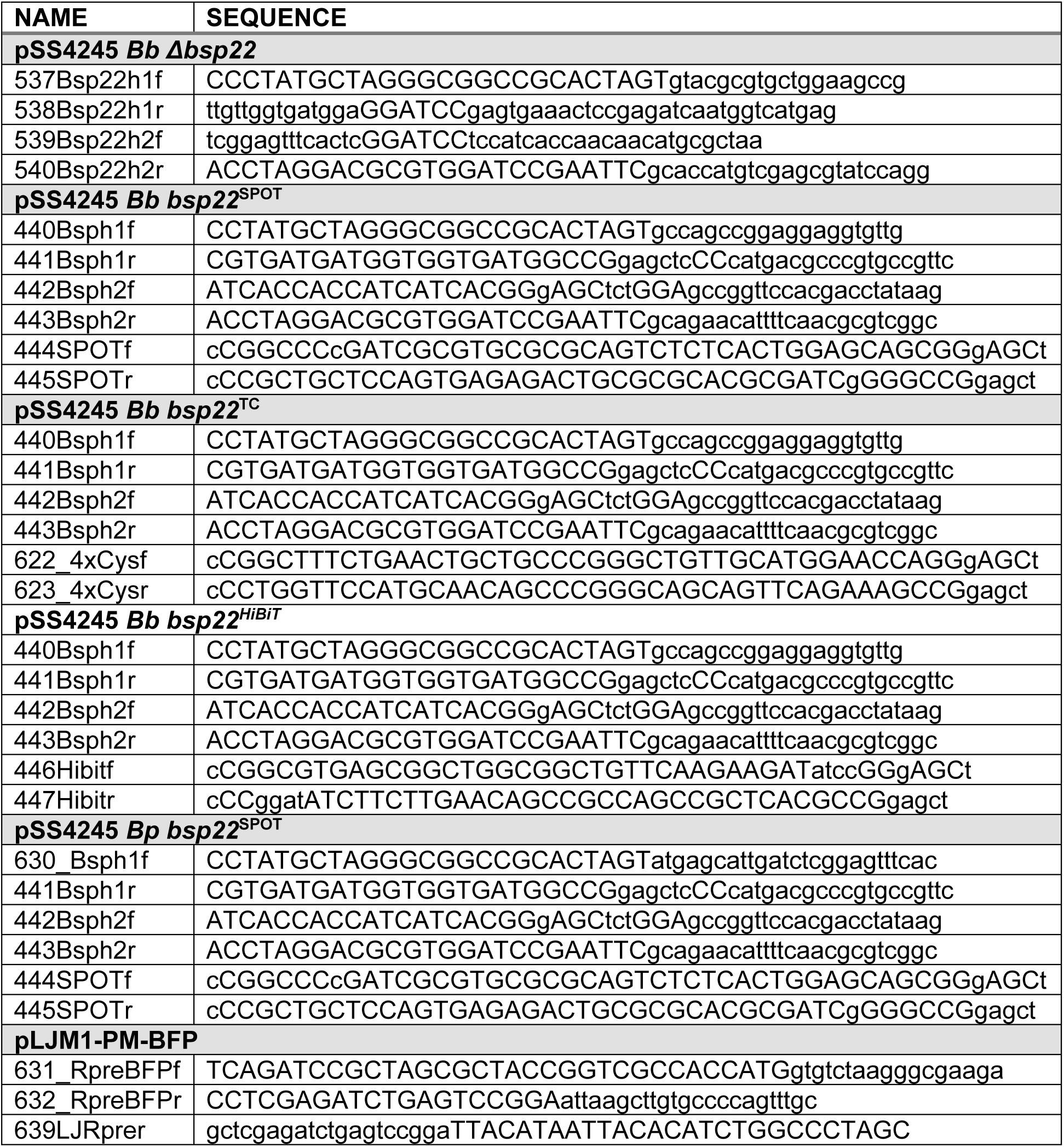
LIST OF OLIGONUCLEOTIDES USED IN THIS STUDY. Oligonucleotides are listed according to the constructed plasmid.

